# Periscope Proteins are variable length regulators of bacterial cell surface interactions

**DOI:** 10.1101/2020.12.24.424174

**Authors:** Fiona Whelan, Aleix Lafita, James Gilburt, Clément Dégut, Samuel C. Griffiths, Huw T. Jenkins, Alexander N. St John, Emanuele Paci, James W.B. Moir, Michael J. Plevin, Christoph G. Baumann, Alex Bateman, Jennifer R. Potts

**Affiliations:** Department of Biology, The University of York, York YO10 5DD, UK; European Molecular Biology Laboratory, European Bioinformatics Institute (EMBL-EBI), Wellcome Genome Campus, CB10 1SD, UK; Department of Chemistry, University of York, York YO10 5DD, UK; Astbury Centre for Structural Molecular Biology, The University of Leeds, Leeds, LS2 9JT, UK

## Abstract

Changes at the cell surface enable bacteria to survive in dynamic environments, such as diverse niches of the human host. Here, we reveal “Periscope Proteins” as a widespread mechanism of bacterial surface alteration mediated through protein length variation. Tandem arrays of highly similar folded domains can form an elongated rod-like structure; thus variation in the number of domains determines how far an N-terminal host ligand binding domain projects from the cell surface. Supported by newly-available long-read genome sequencing data, we propose this new class could contain over 50 distinct proteins, including those implicated in host colonisation and biofilm formation by human pathogens. In large multi-domain proteins, sequence divergence between adjacent domains appears to reduce inter-domain misfolding. Periscope Proteins break this “rule”, suggesting their length variability plays an important role in regulating bacterial interactions with host surfaces, other bacteria and the immune system.

## Introduction

Bacteria encounter complex and dynamic environments, including within human hosts, and have thus evolved various mechanisms that enable a rapid response for survival within, and exploitation of, new conditions. In addition to classical control by regulation of gene expression, bacteria exploit mechanisms that give rise to random variation to facilitate adaptation (e.g. phase and antigenic variation (*1*)). In Gram-positive and Gram-negative human pathogens DNA inversions (*2*, (*3*), homologous recombination (*4*), DNA methylation (*1*) and promoter sequence polymorphisms (*5*) govern changes in bacterial surface components including capsular polysaccharide and protein adhesins, which can impact on bacterial survival and virulence in the host (*1*, (*6*). Many of these mechanisms are very well studied and widespread across bacteria.

A less well-studied mechanism is length variation in bacterial surface proteins. Variability in the number of sequence repeats in the Rib domain (*7*)-containing proteins on the surface of Group B streptococci has been linked to pathogenicity and immune evasion (*8*). The repetitive regions of the *Staphylococcus aureus* surface protein G (SasG) (*9*) and *Staphylococcus epidermidis* SasG homologue, Aap (*10*) also demonstrate sequence repeat number variability. In SasG this variability regulates ligand binding by other bacterial proteins *in vitro (11)* in a process that has been proposed to enable bacterial dissemination in the host. Variations in repeat number have also been noted in the biofilm forming proteins Esp from *Enterococcus faecalis* (*12*) and, more recently, CdrA from *Pseudomonas aeruiginosa (13)*. High DNA sequence identity in the genes that encode these proteins is likely to facilitate intragenic recombination events that would lead to repeat number variation (*14*) and, in turn, to protein sequence repetition. However, such sequence repetition is usually highly disfavoured in large multi-domain proteins (*15*), so its existence in these bacterial surface proteins suggests that protein length variation provides an evolutionary benefit. SasG, Aap and Rib contain N-terminal host ligand binding domains and C-terminal wall attachment motifs, thus our recent demonstration that the repetitive regions of both SasG (*16*) and Rib (*17*) form unusual highly elongated rods suggests that host-colonisation domains will be projected differing distances from the bacterial surface.

Here we show that repeat number variation in predicted bacterial surface proteins is more widespread and we characterise a third rod-like repetitive region in the *S. gordonii* protein (Sgo_0707) formed by tandem array of novel ‘SHIRT’ domains. Thus, we propose a new, and growing, class of “Periscope Proteins”, in which long, highly similar DNA repeats facilitate expression of surface protein stalks of variable length. This mechanism could enable changes in response to selection pressures and confer key advantages to the organism that include evasion of the host immune system (*8*) and regulation of surface interactions (*11*) involved in biofilm formation and host colonisation.

## Results

### Defining the structural repeats of Sgo_0707 from *Streptococcus gordonii*

Having revealed the unusual repetitive, rod-like characteristics of both SasG (*16*) and Rib (*17*) in our previous studies, we used bioinformatic approaches to search for other cell-wall attached bacterial proteins with related organisation. A8AW49 (herein Sgo_0707) encoded by the gene *Sgo_0707* from *S. gordonii* (Fig. 1A) has a C-terminal wall attachment motif, homologues with repeat number variation, and a structurally defined two-domain N-terminus (N1-N2; residues 36–458, PDB: 4igb), that is proposed to be involved in collagen binding (*18*). As the repeats had no Pfam definition we called the putative domain “SHIRT” (Streptococcal High Identity Repeats in Tandem; Fig. 1B). *S. gordonii* is a member of the *S. sanguinis* group of viridans streptococci (*19*) and is a common colonizer of the oral cavity. It is a pioneer organism in the establishment of dental plaque (*20*) and also implicated in infective endocarditis (*21*).

**Fig. 1.**
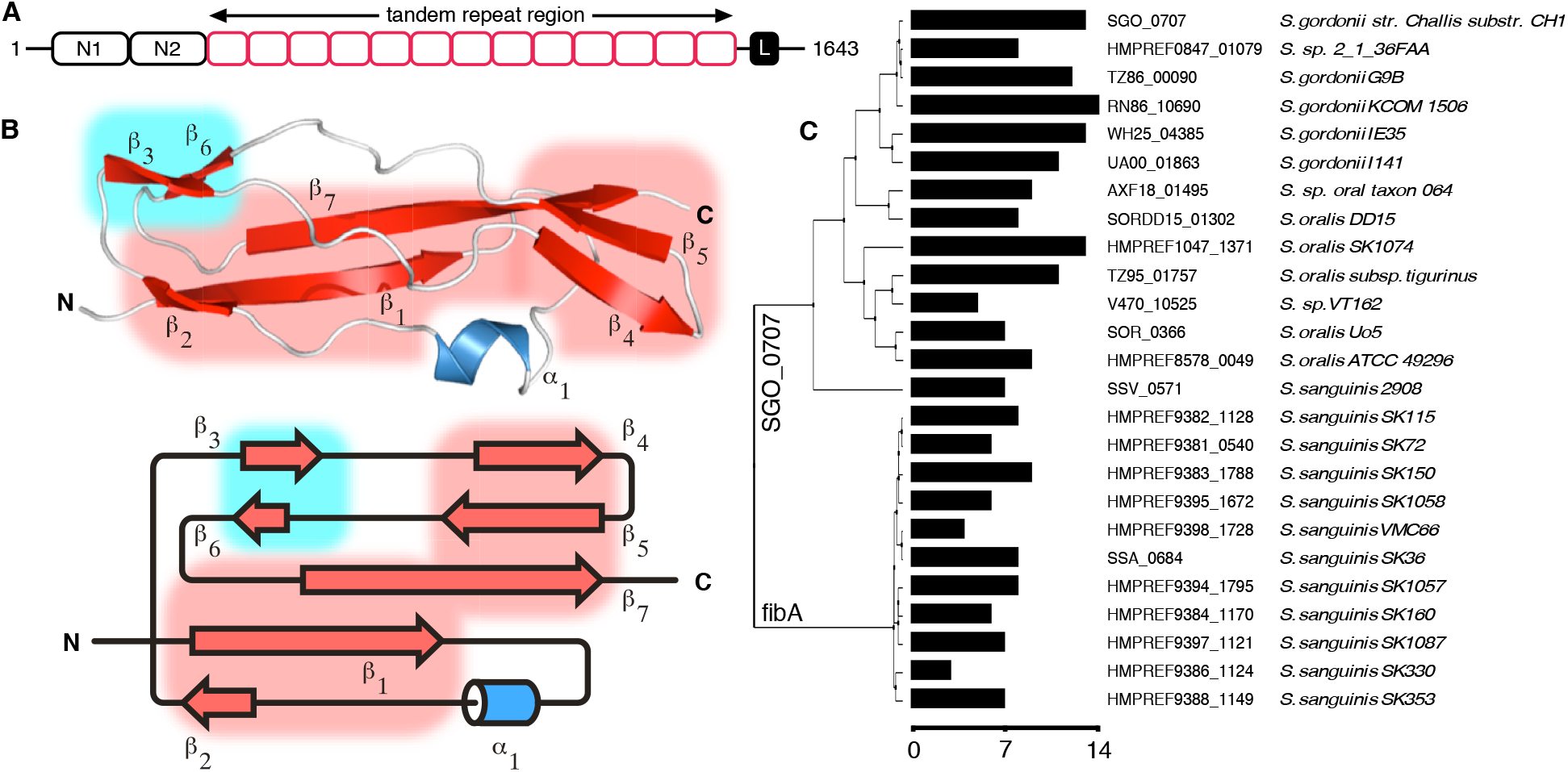
Close homologs of A8AW49 contain variable numbers of repeats that each form the novel ‘SHIRT’ fold. **(A)** Schematic of Sgo_0707 showing the N-terminal adhesin domains N1 and N2, tandem repeat (SHIRT) domains (red) and C-terminal LPNTG cell-wall crosslinking motif (black box ‘L’). **(B)** Structure of Sgo_R10 and topology; sheets S1 (red) and S2 (blue) are highlighted in boxes**. (C)** A phylogenetic tree of Sgo_0707 homologs (>70% identity) identified using PHMMER (*59*) from ENSEMBL bacterial genomes using the N-terminal domain sequence (residue range 1-450 for Sgo_0707) and containing the LPXTG cell wall anchor motif at the C-termini. The number of SHIRT domain repeats in the stalk of each protein is shown as a bar plot with scale bar below.

Defining the structural, rather than sequence, repeat boundaries in repetitive bacterial proteins is challenging. For Sgo_0707 the T-REKS server (*22*) predicts a repeat frame of 460–543 and 13 repeats of 84–90 residues. A construct based on the second repeat (residues 544–627; ΔN-Sgo_R2) was folded (fig. S1A) and solved to 0.95 Å resolution using X-ray crystallography (fig. S1A inset and table S1) utilising *ab initio* molecular replacement (MR) with ideal fragments (*23*). Based on the significant truncation of the N-terminal β-strand (fig. S1A inset), we hypothesised that shifting the frame of the repeat by seven residues towards the N-terminus of Sgo_0707 would complete the fold. Sgo_R3 (residues 621–705) and Sgo_R10 (residues 1211–1299) based on this new definition were thus expressed and purified. They were found to have significantly higher melting temperatures (*T*_m_) than the N-terminally truncated Sgo_R2 (ΔN-Sgo_R2, 55.9°C; Sgo_R3, 75.7°C; Sgo_R10, 75.9°C; fig. S1B).

The structure of Sgo_R10 (Fig. 1B) was solved at 0.82 Å resolution using MR with the structure of ΔN-Sgo_R2 for phasing; the data collection and refinement statistics are summarised in Table 1. The model confirms that SHIRT has a novel α/β fold organized around a single α-helix and two distinct β-sheets (Fig. 1B). Fig. 1A shows a schematic of Sgo_0707 based on the structural boundaries of the repeats; fig. S1C shows the high level of protein sequence identity (82–100%) between adjacent SHIRT domains. Comparison of Sgo_0707 genes from different bacterial strains, including the homologous protein fibA from *S. sanguinis*, shows a high variability in the number of SHIRT domain repeats forming the stalk, ranging from 3 to 14 copies (Fig. 1C). SHIRT domains are found in many other proteins, often in tandem array (fig. S2).

### Tandemly-arrayed Sgo_0707 SHIRT domains form an extended rod-like structure

A tandem domain construct (Sgo_R3-4; residues 621—789) was crystallised and the structure solved via MR using the ΔN-Sgo_R2 model; data collection and refinement statistics are summarised in Table 1. The structure (Fig. 2A) reveals two complete domains with a very short (Pro-Ala-Pro) linker (fig. S1C); the structure of Sgo_R3-4 is ordered throughout (residues 623–789). Each domain adopts the SHIRT fold and the inter-domain interface is limited; this was confirmed by comparing the *T*_m_ of Sgo_R3 (75.7°C) and Sgo_R3-4 (76.6°C; fig. S3). The similarity of unfolding curves for single and double SHIRT constructs suggests that the two domains in the tandem construct unfold independently. Small angle X-ray scattering (SAXS) analysis substantiates the anisotropic head-to-tail domain arrangement in solution (fig. S4A). Notably, there is a significant twist between domains when viewed along the long axis of the molecule.

**Fig. 2.**
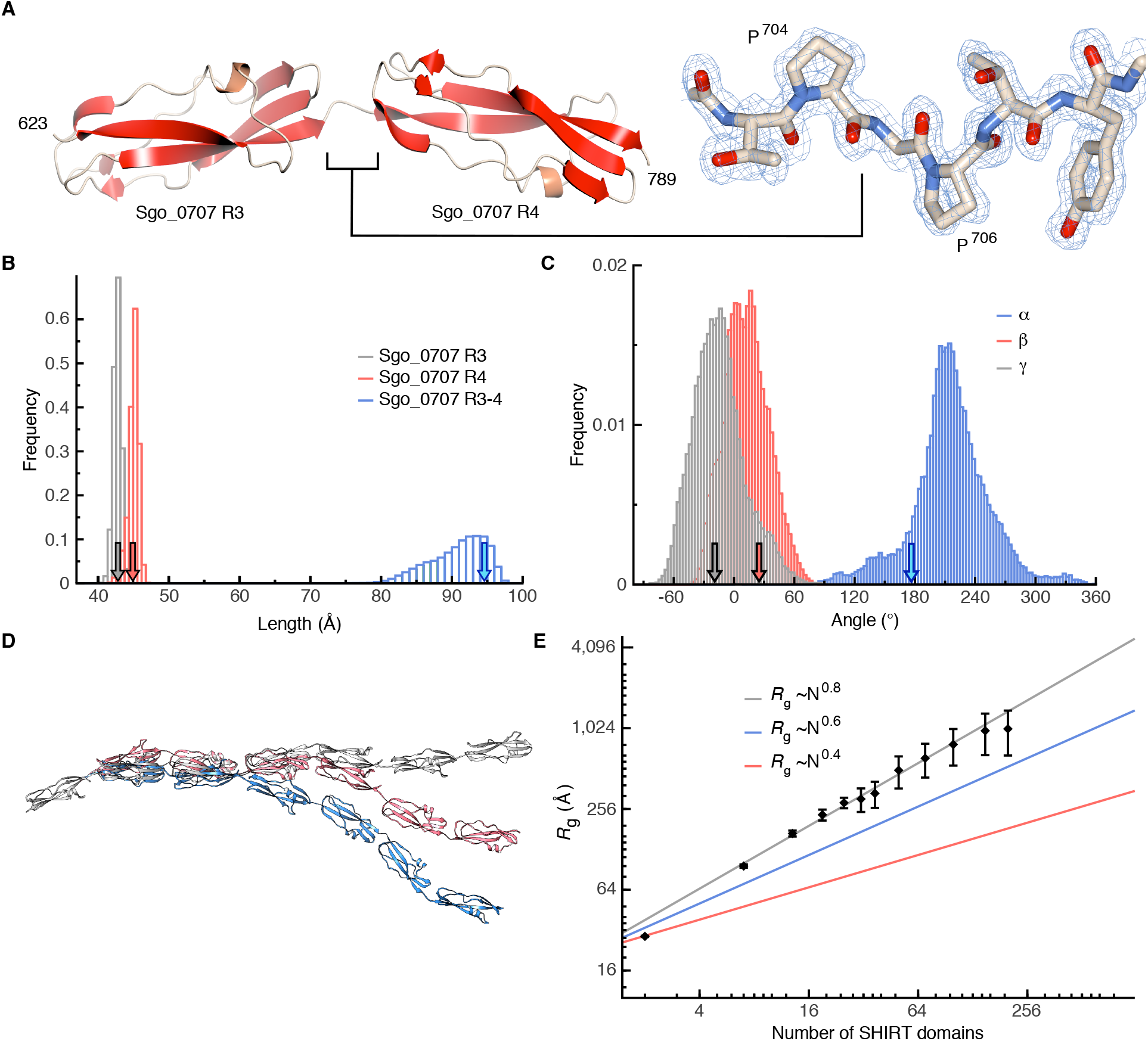
The tandem SHIRT repeat Sgo_R3-4 adopts an anisotropic (head-to-tail) structure with limited inter-domain “bend” but significant “twist”. **(A)** The structure of Sgo_R3-4 (left) and connecting short, well-ordered Pro-Ala-Pro inter-domain linker (electron density 2mF_o_-DF_c_ map contoured at 1σ (blue chickenwire, right); Ala β-methyl group not visible in this orientation). **(B)** Frequency of domain lengths and **(C)** inter-domain angles (fig. S5A) over a 0.8 μs all-atom, fully solvated molecular dynamics simulation of Sgo_R3-4 at 303 K. The ends of the domains Sgo_R3, Sgo_R4 and Sgo_R3-4 and linker are identified by the Cα atoms of residues T623–P704, T707–A789 and T627–A789, respectively; arrows show the value of the distances in the crystal structure. **(D)** Models of 7 tandemly-arrayed domains based on the α, β, γ angles in c. **(E)** Scaling of *R*_g_ of simulated model SHIRT constructs with increasing number of domains (or, equivalently, amino acids) compared to denatured and native proteins (approximated by blue and red lines, respectively). Error bars are standard deviations over 100 models generated.

Molecular dynamics (MD) simulations of the Sgo_R3-4 construct show that individual domains are particularly stable (RMSD <1.5 Å for Cα atoms) over the length of the trajectory (0.8 μs); their individual length is conserved during the simulation and the length of Sgo_R3-4 fluctuates only moderately around 97 Å (Fig. 2B). The distributions of α, β, γ inter-domain (*17*) angles (fig. S5A) observed in the simulations of the Sgo_R3-4 construct (Fig. 2C) were used to generate models of longer constructs (Fig. 2D). The radius of gyration (*R*_g_) of the simulated constructs increases following the relation *R*_g_ ∝ *N*_v_, where *v* is the Flory exponent and describes the increase in size of a polymer (protein) made of *N* monomers (amino acids). Such an exponent is ~0.6 for denatured proteins and ~0.4 for folded ones (*24*). Polymers formed of sequential SHIRT domains are highly extended; *R*_g_ scales with the number of domains (or equivalently, amino acids) with an exponent *v* ~0.8 (Fig. 2E), which is remarkable given the width of the distribution of the angles describing the mutual orientation of adjacent domains (Fig. 2C).

To assess the elongation of Sgo_0707 in solution, we collected SAXS data for constructs comprising two (Sgo_R3-4) and seven (Sgo_R2-8) tandemly-arrayed SHIRT domains (Fig. 3 and fig. S4). Both Sgo_R3-4 and Sgo_R2-8 are monomeric in solution, eluting as monodisperse peaks from size exclusion chromatography (SEC) columns (fig. S4, A and B, inset). Both the crystal structure of Sgo_R3-4 (Fig. 2A) and an elongated model for Sgo_R2-8 (fig. S4B) are consistent with the SAXS data measured in solution (model:data fits of χ^2^=1.1 and χ^2^=1.2, respectively; fig. S4, A and B and Materials and Methods). Analysis of the data for Sgo_R3-4 and Sgo_R2-8 results in equal Porod exponents (1.2, Fig. 3A), as well as similar radii of gyration of a cross-section (*R*_g_^c^) values (R3-4 = 6.4±0.0 Å; R2-8 = 7.0±0.0 Å; Fig. 3, B and C). Consistent with the models (Fig. 2D), the *D*_max_ determined using SAXS scales with the number of domains; Sgo_R3-4 exhibits a *D*_max_ of 107 Å and Sgo_R2-8 has a *D*_max_ of 371 Å (fig. S4, A and B). Therefore, whilst displaying a much larger intra-particle maximum dimension, the Sgo_R2-8 construct has a comparable shape and *R*_g_^c^ to Sgo_R3-4.

**Fig. 3.**
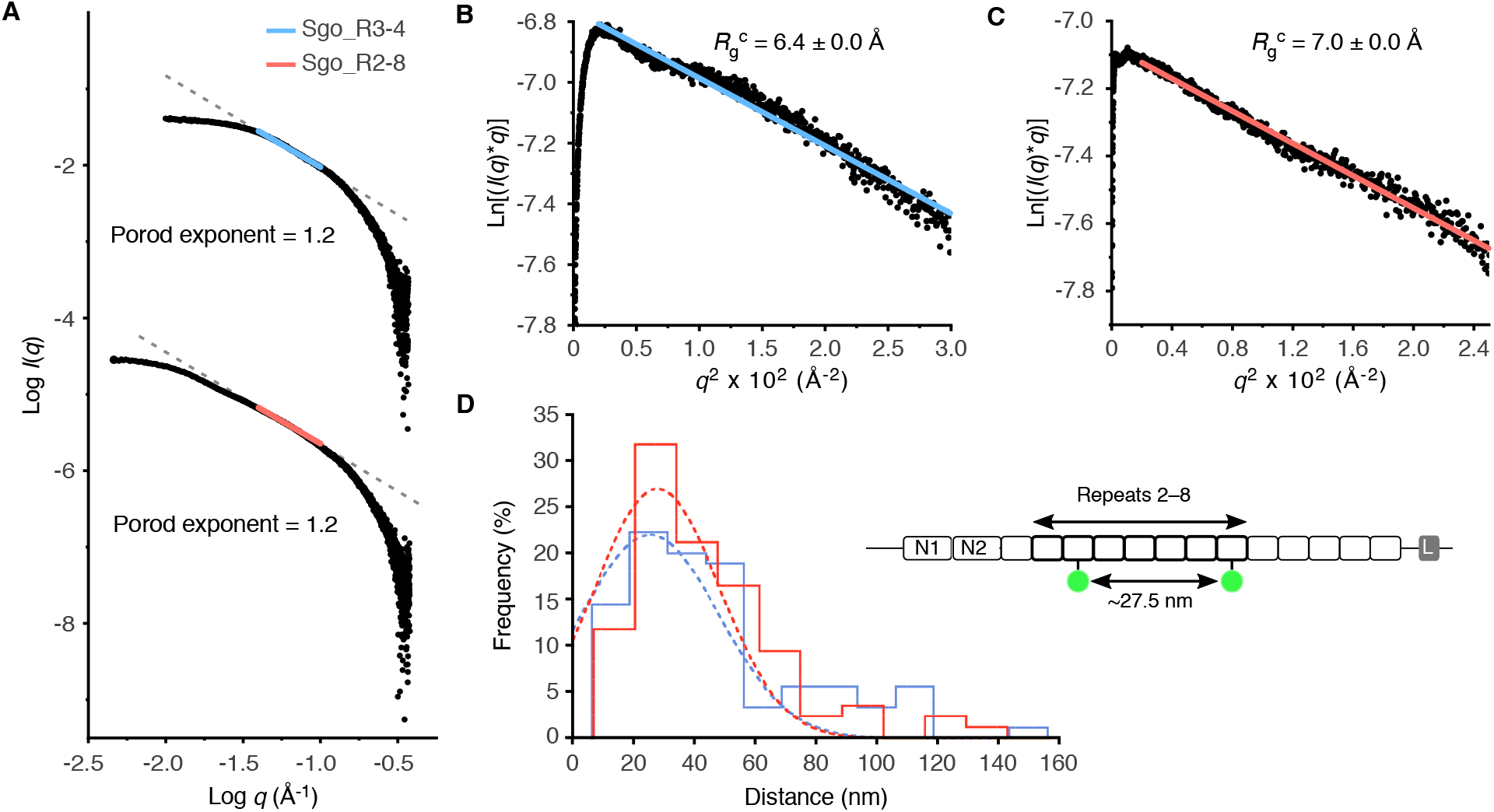
Tandemly-arrayed SHIRT domains form a highly elongated rod. **(A)** Porod plots from SAXS analysis of Sgo_R3-4 (top) and Sgo_R2-8 (bottom). Porod exponents were calculated from the negative slope of the linear region of the data (R3-4, blue, *R*^2^ = 1.0; R2-8, red, *R*^2^ = 1.0). **(B)** Modified Guinier region for Sgo_R3-4 with cross-sectional radius of gyration (*R*_g_^c^, annotated) calculated from fitted region (blue line, *R*^2^ = 0.98, *q*\**R*_g_ limits = 0.3–1.1). **(C)** Modified Guinier region for Sgo_R2-8 with *R*_g_^c^ calculated from fitted region (red line, *R*^2^ = 0.99, *q*\**R*_g_ limits = 0.3–1.0. **(D)** SHRImP-TIRFM determination of the inter-dye distances for AF488-labelled Sgo_R2-8^S666C/S1086C^ (schematic inset) immobilised on 2 μg/mL (blue) or 20 μg/mL (red) poly-D-lysine treated quartz surfaces. Colour-coded, dashed lines indicate Gaussian fits to the histograms for the 2 μg/mL (*n* = 90, *R*^2^ = 0.86) and 20 μg/mL (*n* = 85, *R*^2^ = 0.90) poly-D-lysine-treated quartz surfaces (mean ± standard error = 25.5 ± 2.39 nm and 27.8 ± 2.21 nm, respectively).

In our previous study (*17*), we observed elongation in 2-domain Rib constructs with rotation of angle α, whilst angles β and γ were smaller. We conducted the same analysis for Sgo_0707 constructs, based on fitting to our experimental SAXS data (fig. S5). As with Rib, MD simulations of the 2-domain Sgo_R3-4 construct show that a range of α angles give good fits to the observed SAXS data, whilst β and γ are restricted to a narrow range, consistent with an elongated conformation (fig. S5B). For fitting of MD simulations of the 7-domain Sgo_R2-8 construct to SAXS data, we observed that longer end-to-end distances improved the quality of the fit (fig. S5C). Taken together, these data are consistent with multiple tandemly-arrayed SHIRT domains from Sgo_0707 behaving as an elongated, rod-like particle.

To further assess the elongation of multi-domain SHIRT constructs, we used a high-resolution single-molecule technique (SHRImP (*16*)) to measure the intramolecular distance between two Alexa Fluor 488 (AF488) dyes covalently attached to cysteine residues engineered at specific sites in Sgo_R2-8 (S666C and S1086C). If the extended inter-domain topology observed in the two-domain construct is maintained, a distance of 24.1 nm between the mean dye positions (fig. S6A) is predicted by using the distance between the two cysteine residues (Fig. 3D inset) and simulating the increased volume accessible to each dye due to the chemical linker (*25*). AF488-labelled Sgo_R2-8^S666C/S1086C^ was imaged using total internal reflection fluorescence microscopy (TIRFM) using two different poly-D-lysine concentrations to coat the slides. Both surface treatments effectively immobilised AF488-labelled Sgo_R2-8^S666C/S1086C^ and each SHRImP-TIRFM histogram had a single peak consistent with monodispersity. The measured inter-dye distances were 25.5 nm and 27.8 nm (Fig. 3D) with 2 μg/mL and 20 μg/mL poly-D-lysine concentration, respectively, and increased to 29.8 nm and 33.0 nm, (fig. S6B), respectively, when measurements were performed in the presence of 100-fold molar excess of unlabelled Sgo_R2-8 (termed ‘blocking’ protein). This implies the rod-like conformation of Sgo_R2-8 protein is malleable, adopting, compared to solution, a slightly more elongated state when on a surface which is accentuated (Δx = 4-5 nm) at high protein concentrations that presumably favour lateral protein-protein interactions.

### Large-scale identification of potential Periscope Proteins in bacterial genomes

We call SasG, Rib and Sgo_0707 ‘Periscope Proteins’ due to their having variable length stalk-like regions that define the distance that the N-terminal functional domain projects from the bacterial surface. We conducted an unbiased identification of other potential Periscope Proteins from the NCTC3000 bacterial genomes, an ongoing project lead by the Wellcome Sanger Institute (UK) that provides high quality annotated genome assemblies for 3,000 bacterial strains from Public Health England's National Collection of Type Cultures (NCTC) and sequencing using the long-read PacBio technology (https://www.sanger.ac.uk/resources/downloads/bacteria/nctc/). Using long-reads spanning whole genes overcomes the challenges in the assembly of repetitive genes in bacterial genomes (*26*), providing reliable numbers of repeats in the assembled genes and associated proteins, and allowing the identification of length variability in Periscope Proteins. We identified 1,576 proteins containing long and highly identical repeats (i.e. similar to those forming stalks in Periscope Proteins), out of a total of 2.5 million proteins in the NCTC3000 dataset, from different species including both Gram-positive and Gram-negative bacteria. Clustering of the repeating sequences reveals they mostly fold into globular domains, as in known Periscope Proteins, and most of them are classified into existing Pfam families (Fig. 4).

**Fig. 4.**
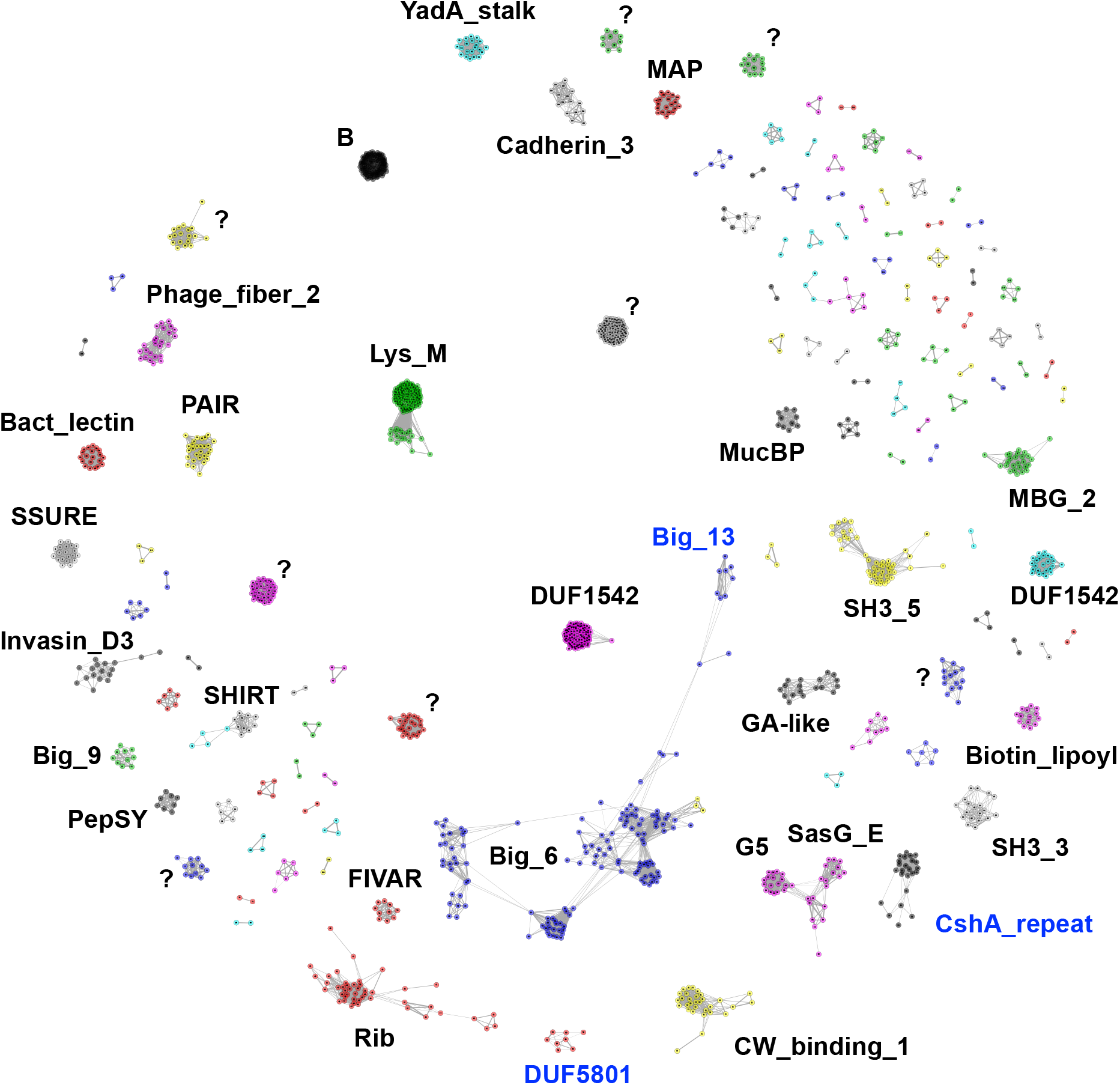
Sequence clustering of long tandem highly identical repeats identified across proteins of the NCTC3000 genomes. The largest clusters are annotated using Pfam. Clusters with unknown Pfam classification are marked with “?”. New protein domain families built from sequence clusters into Pfam are highlighted in blue.

We further clustered full-length repetitive proteins and identified a total of 180 unique groups, 84 of which exhibit repeat number variation (Supplementary Data 1). Manual inspection confirmed that 56 of these groups are examples of Periscope Proteins, 30 of which could be assigned to recognisable gene names. We retrieved known Periscope Proteins, such as SasG, Rib and Sgo_0707 (clusters 4, 25 and 26 respectively), as well as novel ones, for example CdrA from *Pseudomonas aeruginosa (27)* which has been reported recently to exhibit variability in the size of the repeat region (*13*), CshA from *S. gordonii (28)*, and SasY from *S. aureus* (Fig. 5). The majority of confirmed Periscope Proteins (40/56) have at least one Gene Ontology (GO) location term assigned, all of them associating to the cell-wall or bacterial membrane.

**Fig. 5.**
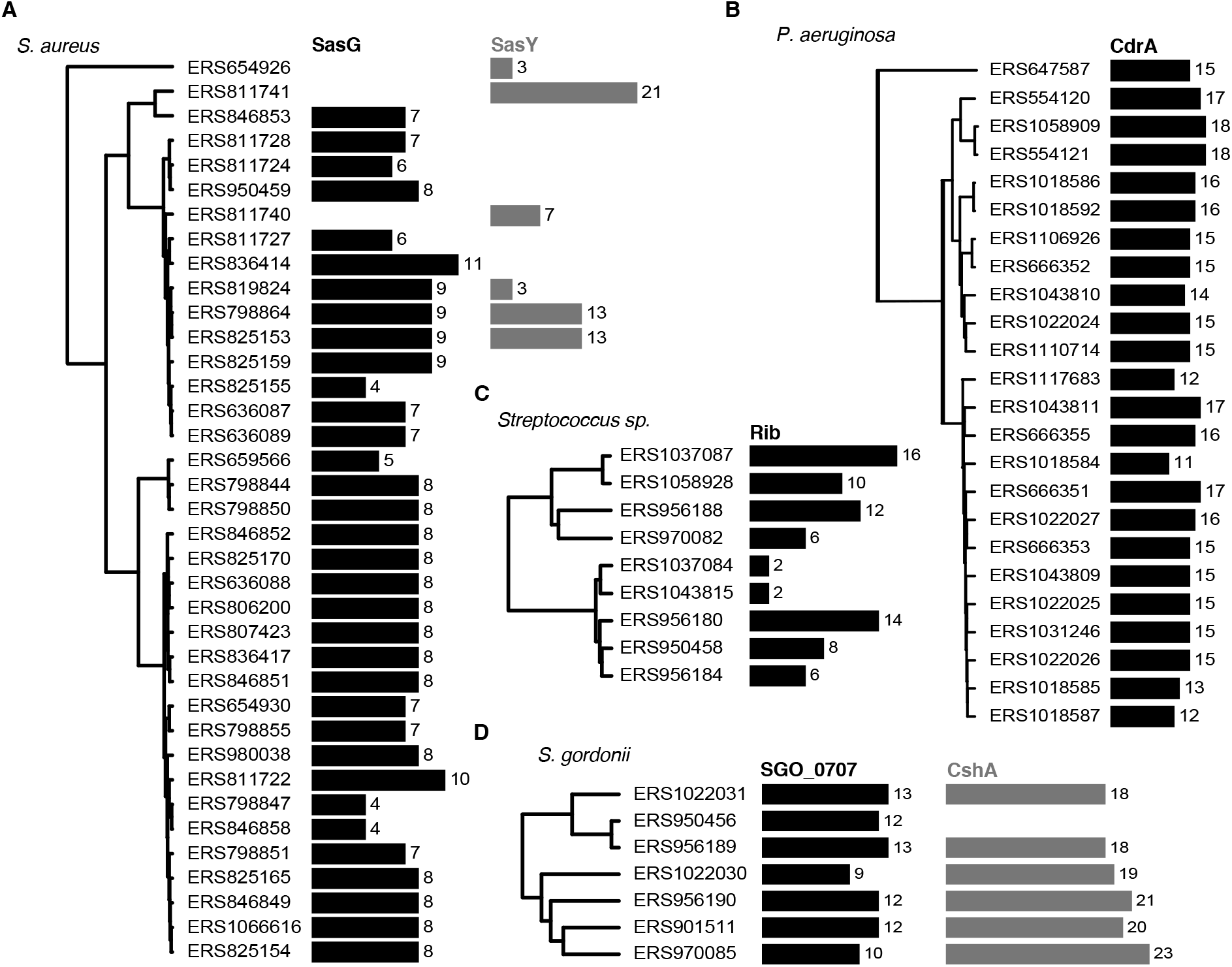
Variation of stalk region repeat numbers in Periscope Proteins. Phylogenetic trees of **(A)** *Staphylococcus aureus* **(B)** *Pseudomonas aeruginosa* **(C)** various streptococcal species including *S. agalactiae* and *S. pyogenes* and **(D)** *S. gordonii* genomes in the NCTC3000 collection, mapped to the number of repeats in stalk regions in Periscope genes; respectively, SasG with G5/E repeats, SasY with Big_6 domain repeats, CdrA with MBG_2 domain repeats, surface protein Rib with Rib domain repeats, Sgo_0707 homolog containing SHIRT domain repeats, and surface adhesin CshA with ~100-residue globular domain repeats of a new Pfam family (CshA_repeat).

We further find that the number of repeats per gene can be large (>40 in one case (*29*); Fig. 6), spanning thousands of amino acids, and that variation in the number of stalk repeats can be extreme, increasing the length of the protein by an order of magnitude in some cases. Importantly, we found the most extreme repeat numbers and length variation in proteins with the highest repeat identity (Fig. 6). Furthermore, as shown in Fig. 5, the number of repeats changes drastically even between similar bacterial strains, suggesting a rapid evolutionary rate of repeat number change in Periscope Proteins.

**Fig. 6.**
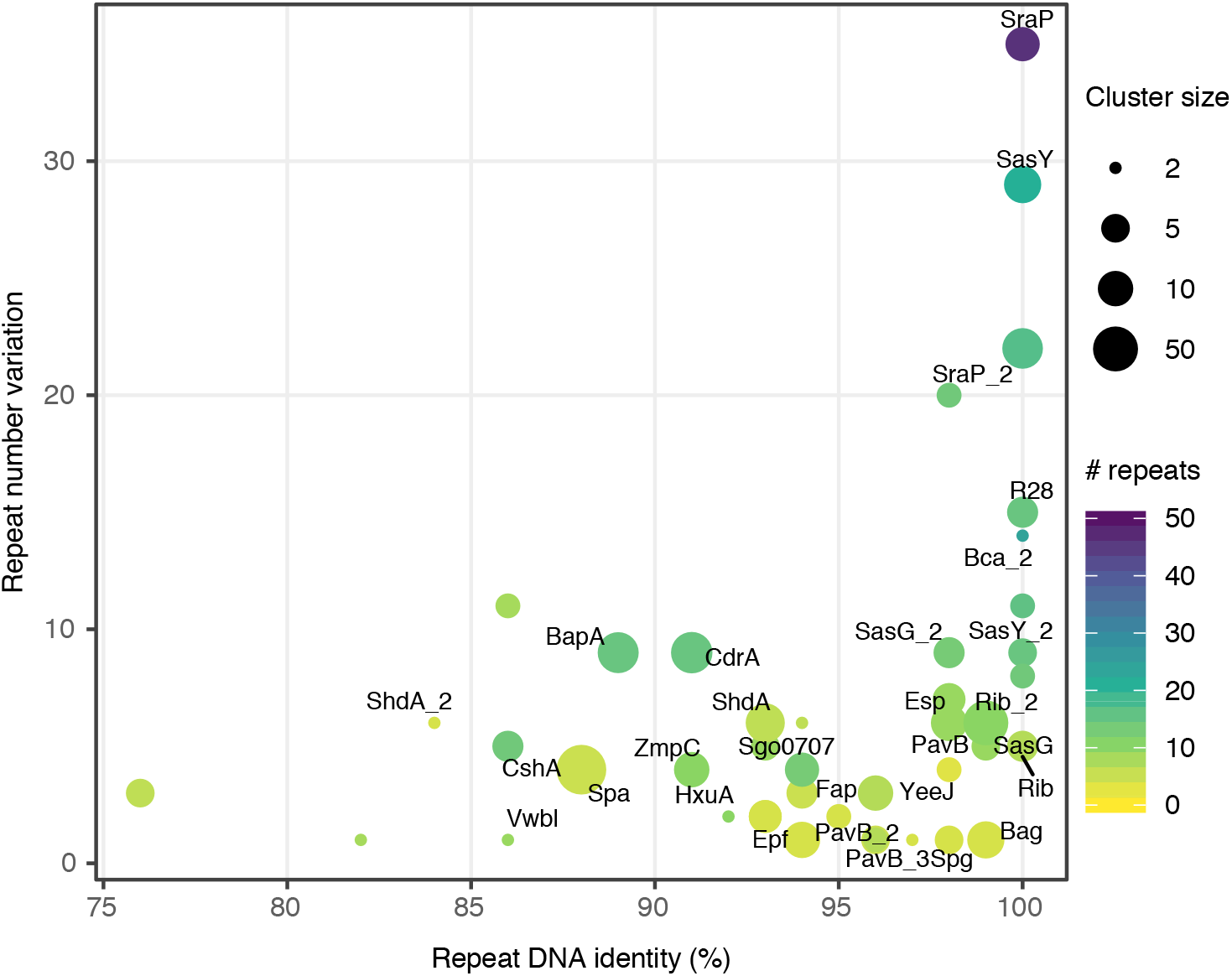
Repeat number variation in Periscope Proteins as a function of repeat sequence identity. The sequence identity of tandem repeats in the genome plotted against the variation in repeat number observed for each Periscope Protein cluster. The repeat number variation is calculated as the difference between the maximum and minimum observed repeat numbers in proteins within each cluster. The maximum number of repeats is shown as a viridis color scale. The repeat DNA identity is calculated as the maximum repeat sequence identity across proteins in each cluster. The number of proteins in each cluster is shown as point size. Names for the most relevant Periscope Protein clusters are shown as labels.

## Discussion

The work presented in this study identifies a new class of bacterial surface proteins that, through length variation, are likely to modulate surface interactions. Through biophysical analysis of three unique examples (SasG (*16*), and Rib (*17*) previously and Sgo_0707 in this study) and bioinformatic analyses (this study), we show this class is characterised by a highly repetitive central region with a variable number of domain-forming repeats in tandem. The domains in tandem can form an elongated rod that typically projects a functional domain distal to the cell surface. To classify this diverse family of proteins with a memorable analogy, we call them ‘Periscope Proteins’. We propose that high sequence similarity at the DNA level enables intragenic recombination events (and thus loss or gain of repeats) with selection pressures resulting in enrichment of bacteria with shorter or longer proteins. Periscope Proteins include proteins implicated in biofilm formation in both Gram-positive and Gram-negative bacteria, for example SasY and related proteins in *S. epidermidis*, *S. xylosus*, *S. capitis, S. simiae* and *S. warneri*, the *Enterococcus faecalis* protein Esp (*12*) and related proteins, and CdrA from *Pseudomonas aeruginosa (13)*. We expect the total number of Periscope Proteins across bacterial species to be much larger and more widespread than reported in this study. Our knowledge of Periscope Proteins will increase as more long-read sequence data for a diverse set of strains not included in the NCTC3000 collection become available.

Large multi-domain proteins usually have <50% sequence identity between adjacent domains (*15*) which has been proposed as an evolutionary strategy to minimise inter-domain misfolding (*30*, (*31*, (*32*). The existence of Periscope Proteins, which contain tandemly-arrayed, structured domains with high sequence similarity, confounds this observation. Considering this potential “misfolding problem”, the existence of Periscope Proteins suggests that length variation (driven by the requirement for high identity at the DNA level, and thus high protein similarity) is functionally important and confers a significant advantage to the organism. Simply expressing a Periscope Protein (an extreme form of length variation) results in changes in ligand binding by other bacterial surface proteins through spatial competition. For example, expression of Pls (*33*) and SasG significantly inhibits binding of *S. aureus* to the human plasma proteins fibronectin and fibrinogen, respectively; the latter resulting in altered bacterial colony morphology (*34*). In both cases it is suggested that the elongated Periscope Protein is blocking interactions of the host protein with bacterial proteins closer to the cell surface. There is also more direct evidence for the role of length *variation* in Periscope Proteins under selection pressure. For example, in mice immunised with anti-serum raised against the nine-repeat alpha C, GBS expressing alpha C with one Rib repeat were 100-fold more pathogenic than GBS expressing alpha C with nine Rib repeats (*35*). It was proposed that the shorter protein is not recognised because it is less exposed. When SasG was expressed on *S. aureus* with differing numbers of repeats, only the longer variants blocked binding of other bacterial surface proteins to their target (*11*). These authors noted that the ability of a bacterium to detach from a ligand could be important for dissemination in the host. Finally, deletion of CdrA had the most deleterious effect on biofilm formation for *P. aeruginosa* strains expressing the longest CdrA variants (*13*). Rib repeat number variation within a single strain can be observed within 24 hours following inoculation of a mouse with group B streptococci (*36*), suggesting the potential for dynamic regulation by Periscope Proteins on a physiologically relevant timescale. In summary, surface variation through the Periscope Protein class appears to be a highly distinctive way to alter a variety of interactions linked to colonisation and infection.

## Materials and Methods

### Cloning

The *E. coli* codon-optimised coding sequence for *S. gordonii* strain Challis (substrain DL1) Sgo_0707 residues 544–795 (UniProt: A8AW49) was synthesized (Genewiz; Supplementary Table 2) and the truncated single repeat ΔN-Sgo_R2 (amino acids 544–627), single repeats Sgo_R3 (aa 621–705) and Sgo_R10 (aa 1211–1299) (synthesized by Eurofins Genomics), and tandem repeat Sgo_R3-4 (aa 621– 789) were cloned downstream of a hexahistidine tag and 3C protease specific linker by the In-fusion method (Clontech; primer sequences listed in Supplementary Table 1). DNA coding for the 7-repeat construct (Sgo_R2-8) comprising the 2^nd^ to 8^th^ SHIRT repeats (aa 537–1125) was synthesised (Eurofins Genomics), and inserted by homologous recombination into a modified pBAD vector (pBADcLIC2005) generating a coding sequence comprising GGGFA-Sgo_R2-8-His10. Site-directed mutagenesis was used to introduce cysteine mutations in the Sgo_R2-8 construct for fluorescent dye modification at positions S666 and S1086 (Sgo_R2-8^S666C/S1086C^; mutagenesis primer sequences listed in Supplementary Table 1).

### Expression and purification

For ΔN-Sgo_R2, Sgo_R3 and Sgo_R3-4, expression was induced in *E. coli* BL21 (DE3) cells in log phase growth with the addition of 0.1 mM isopropyl β-D-1-thiogalactopyranoside (IPTG) and subsequently incubated with shaking at 20°C for 20 h. Cells were harvested by centrifugation, resuspended in 20 mM Tris.HCl, 150 mM NaCl, 20 mM imidazole, pH 7.5, and lysed by sonication. Soluble protein was purified by standard nickel affinity chromatography methods, 3C protease was added at a ratio of 1:100 (w/w) to remove the hexahistidine tag, and proteolysed material isolated by a second nickel affinity chromatography purification. Sgo_R10 and Sgo_R3-4 were further purified by preparative SEC using a Superdex 75 16/600 column (GE Healthcare) equilibrated in 20 mM Tris.HCl, 150 mM NaCl, pH 7.5 (Sgo_R10) and pH 8 (Sgo_R3-4). Sgo_R2-8 constructs were transformed into Rosetta-2 E. coli (DE3) cells and grown in LB-Miller media containing 100 μg/mL ampicillin at 37°C to an OD_600_ of 0.5-0.6. Gene expression was induced by addition of 0.1% (w/v) L-arabinose, followed by incubation at 20°C. Soluble protein was purified as described for Sgo_R10. Protein containing cysteine mutations were purified by affinity chromatography and SEC with the addition of 5 mM β-mercaptoethanol at all steps. Protein samples prepared for SEC-SAXS were dialysed into 20 mM Tris.HCl, 150 mM NaCl, 1 mM EDTA, pH 7.5.

### Protein crystallisation

Proteins were concentrated (ΔN-Sgo_R2 to 48 mg/mL; Sgo_R10 to 30 mg/mL and Sgo_R3-4 to 35.2 mg/mL) by centrifugal filtration with 3 kDa molecular weight cutoff (MWCO) PES (Vivaspin). ΔN-Sgo_R2 crystallised by sitting drop vapour diffusion within 9 months in conditions comprising 0.1 M HEPES pH 7, 2.4 M ammonium sulphate at 291 K. This crystal was passed through 4 M sodium malonate prior to flash cooling. Sgo_R10 crystallised in 5 weeks at 277 K in conditions comprising 2.2 M ammonium sulphate and 150 mM potassium thiocyanate. The crystal was harvested under mineral oil and flash cooled in liquid nitrogen prior to data collection. Sgo_R3-4 crystallised in 4 days in conditions comprising 65% (v/v) 2-methyl-2,4-pentanediol and 0.1 M Tris.HCl pH 8 at 277 K and flash cooled in liquid nitrogen prior to data collection.

### Structure determination

X-ray data were collected at 100 K on the I03 beamline at Diamond Light Source (Didcot, UK). using a Pilatus 3 6M detector. Data were indexed and integrated by XDS (*37*) and scaled and merged by Aimless (*38*). The structure of ΔN-Sgo_R2 was solved by MR with 2 antiparallel 5 residue ideal β-strands using Fragon (*23*) and the model built automatically with ARP/wARP (*39*). Sgo_R3-4 and Sgo_R10 were phased by molecular replacement with Phaser (*40*) using search model ΔN-Sgo_R2. Models were manually-built using Coot (*41*) and refined to completion with REFMAC5 (*42*) for Sgo_R3-4 and PHENIX (*43*) for Sgo_R10 (Table 1). The coordinates and structure factors have been deposited in the protein data bank with accession codes (ΔN-Sgo_R2, 7AVJ; Sgo_R10, 7AVK; Sgo_R3-4, 7AVH). The structures were aligned by secondary structure matching with Superpose (*44*) and cartoons rendered with CCP4mg (*45*) with secondary structure defined by the database of protein secondary structure assignment (DSSP) (*46*).

### Determination of *T*_*m*_

Differential scanning fluorimetry (DSF) was peformed using a Nanotemper Prometheus NT.48 instrument. Protein concentrations were 1 mM and solution conditions were 20 mM Tris.HCl pH 7.5, 150 mM NaCl.

### Small Angle X-ray Scattering (SAXS)

SAXS experiments were performed at beamline B21, Diamond Light Source (Didcot, UK) over a momentum transfer range (*q*) of 0.01 Å^−1^ < *q* < 0.4 Å^−1^. Scattering intensity (*I* vs *q*, where *q* = 4πsinθ/*λ* and 2θ is the scattering angle) was collected using a Pilatus 2M detector, with a beam-to-detector distance of 4014 mm and an incident beam energy of 12.4 keV. Sgo_R3-4 and Sgo_R2-8^S666C/S1086C^ were injected on an inline Shodex KW-203.5 column equilibrated in 20 mM Tris-HCl, 150 mM NaCl, 3 mM KNO3, 5mM β-mercaptoethanol pH 7.5, each at 7.5 mg/mL, and data processing and reduction was performed with Chromixs (*47*). *R*_g_^c^ (Ln[*I*(*q*) vs *q*^2^], and Distance Distribution (*P*(*r*)) plots were calculated with Primus (*48*). The range of useful scattering angles was assessed using Shanum (*49*). SWISS-MODEL (*50*) was used for the generation of a structural model of Sgo_R2-8, using the Sgo_R3-4 crystal structure to generate iterative tandem domains for R3-5, R5-6 and R7-8. Sgo_R2 of the Sgo_R2-8 model was generated based on Sgo_R4 from the Sgo_R3-4 structure. For validation of the Sgo_R3-4 crystal structure, tag residues unresolved in the crystal structure were added using Modeller and all-atom ensembles generated using Allosmod (*50*). In each case, 50 independent pools of 100 models were created, and calculation and fitting of theoretical scattering curves to experimental data was performed using FoXS (*51*). This process was automated using Allosmod-FoXS (*52*). Plots were generated using OriginPro v9.5.5.409 (OriginLab), as were the gradients of the linear regions of double logarithmic plots for the calculation of Porod exponents (*53*).

### Production of fluorescently-labelled protein

As described above, cysteine residues were engineered into the 3^rd^ and 8^th^ repeats of the Sgo_R2-8 construct at positions (S666, S1086), where fluorescence quenching by nearby amino acid residues is minimised. Sgo_R2-8^S666C/S1086C^ (27 μM) was dialysed into 150 mM NaCl, 20 mM Tris.HCl, 1 mM EDTA, pH 7.5, followed by dialysis into 150 mM NaCl, 20 mM Tris.HCl, 1.35 mM tris(2-carboxyethyl)phosphine (TCEP), pH 7.5. The protein was then reacted at 20°C in low light conditions with a 20x molar excess of Alexa Fluor 488 C5 maleimide (ThermoFisher, 540 μM), which was added step-wise over a period of 1.5 h using a 10 mM stock in anhydrous DMSO. The labelling reaction was quenched by adding dithiothreitol (DTT) at 10x molar excess to the maleimide. The protein solution was dialysed into 150 mM NaCl, 20 mM Tris.HCl, 1 mM DTT, pH 7.5 prior to purification by SEC on a S200 30/10 column (Amersham) equilibrated in the same buffer to remove remaining free dye. Purified proteins were stored at −80°C. The labelling efficiency (~2 fluorophores/protein) was estimated from the spectrophotometrically determined concentrations of fluorophore (ε495 _nm_ = 72000 M^−1^ cm^−1^) and protein (ε280 nm = 99475 M^−1^ cm^−1^) after correction for absorption at 280 nm by the fluorophore.

### Sample preparation for SHRImP-TIRF microscopy

A 100 mM Trolox solution was freshly prepared by solubilising 25 mg of Trolox powder (Fluka) in 50 μL methanol, followed by dilution with 850 μL of 0.31 M NaOH solution. All stock buffer solutions were passed through a 0.22 μm pore filter. Adsorption buffer contained 10 mM HEPES, 10 mM NaCl, pH 7.0, 1 mM Trolox, 0.02% (w/v) 5 μm diameter silica beads (Bangs Labs), 8 pM Alexa Fluor 488 (AF488)-labelled Sgo_R2-8^S666C/S1086C^ protein and, in the ‘blocked’ samples only, 800 pM unlabelled Sgo_R2-8 protein. Imaging buffer contained 10 mM HEPES, 10 mM NaCl, pH 7.0, 1 mM Trolox. Poly-D-lysine-coated quartz slides were prepared as described previously (*16*). 1 μM AF488-Sgo_R2-8^S666C/S1086C^ and 200 nM Sgo_R2-8 stock solutions were thawed from −80°C storage and diluted with 20 mM Tris.HCl, 150 mM NaCl (pH 7.5) buffer to the desired concentration before addition to the adsorption buffer at a 25x or 50x dilution. In ‘blocked’ samples, the molar concentration of unlabelled Sgo_R2-8 in the adsorption buffer was maintained at 100x the molar concentration of AF488-Sgo_R2-8^S666C/S1086C^.

In low light conditions, 50 μL of adsorption buffer was distributed along the centre-line of a 2 μg/mL or 20 μg/mL poly-D-lysine-coated quartz slide and then covered with a clean coverslip (No. 1, 22 mm x 64 mm, Menzel-Gläser). A flow chamber was created by sealing the two opposite, short sides of the coverslip with nail varnish. A small amount of imaging buffer was added along the unsealed sides of the flow chamber to prevent it drying out. After 10 min of incubation at room temperature (20-22°C), ~500 μL of imaging buffer was flowed through the chamber created by the slide-silica bead-coverslip sandwich to wash away unbound protein, and the chamber was sealed with nail varnish.

### SHRImP-TIRF microscopy

Fluorescence excitation and detection of AF488 dye emission was achieved using a custom, prism-coupled TIRF microscope as described previously (*16*) with the following modification and increased optical magnification. Quantum dots were not routinely added to the adsorption buffer as an image-focusing aid. Video data (100 frames) were collected using an Evolve 512 (Photometrics) electron-multiplying CCD camera (500 ms exposure) and the pixel size was equivalent to 96 nm in the magnified image.

### Detection and localisation of single fluorophores

Fluorescent spot detection and calculation of inter-AF488 dye distances for each spot that photobleached in two-steps were performed as previously described (*16*). In addition, an eccentricity ratio was calculated for each spot (using the intensity profile in the *x* and *y* directions) to remove events that included weakly surface-adsorbed AF488-labelled protein or partially photobleached clusters of AF488-labelled protein. The intensity profile was calculated for a central region (2 x 10 pixels^2^) along the *x* and *y* axes of each spot image. This image was the sum of the first 5 video frames for each 10×10 pixel^2^ image stack (100 frames). The *x* and *y* intensity profiles were fit with a one-dimensional Gaussian function in MATLAB (MathWorks, Cambridge, UK) and a ratio of the widths for the Gaussian fits (= σ_*y*-axis_ / σ_*x*-axis_) was used to obtain the spot eccentricity. Fluorescent spots with an eccentricity ratio between 0.9 and 1.1 were retained (this filter removed 15–30% of the spots in an experiment). Bin size for the inter-AF488 dye distance histograms was calculated using the Freedman-Diaconis rule (*54*), and a single Gaussian distribution was fit to each histogram in KaleidaGraph (Synergy Software).

### Molecular dynamics simulations

Molecular dynamics simulations were performed starting from the X-ray crystal structure model of Sgo_R3-4. Simulations have been performed using the CHARMM36m (*55*) force field and NAMD (*56*). The protein was energy minimized and solvated in a periodic rectangular box 123 Å x 46 Å x 40 Å needed to guarantee a layer of at least 12 Å solvent around the elongated protein. After a 1 ns equilibration, the systems were simulated at 303K for 0.8 μs. Simulations were performed in the isothermal-isobaric ensemble, where the temperature was kept constant on average through a Langevin thermostat and the pressure was set to 1 atm through an isotropic Langevin piston manostat. For each saved frame of the simulation (one every 2 ns) the positions of the Ca atoms of Sgo_R4 were least square superposed to those of Sgo_R3, which implies a translation and a rotation around the each of the three principal axes of Sgo_R3 (hence applying the reverse transformation to the coordinates of Sgo_R4 the original conformation of that frame is recovered). Constructs with N domains were obtained by taking the original X-ray crystal structure, duplicating the coordinates of Sgo_R4 and applying to them the reverse transformation (using translation and rotations from random frames of the trajectory) N times.

### Periscope protein identification in NCTC3000 collection

A total of 2,579,577 proteins were extracted from 734 annotated bacterial genomes downloaded from the NCTC3000 project website (October 2019). Tandem repeats longer than 50 residues and with at least 80% sequence identity were detected with the T-REKS tool. Repeat sequences were clustered using BLASTp with a bit score threshold of 30 and extracting connected components of the resulting sequence similarity network (SSN). Similarly, proteins containing the long repeats were clustered using BLASTp with a bit score threshold of 100 and additional minimum sequence identity of 90% and coverage of 50% thresholds, by extracting the connected components of the SSN. Proteins were mapped to UniProt (*57*) identifiers (version 2020_04), with their associated Gene Ontology (GO) terms, and to Pfam (*58*) families (version 32.0) using PHMMER. Several new Pfam families were created from repeat sequences during the course of this study, including SHIRT (PF18655), YDG (PF18657), MBG_2 (PF18676), SSSPR-51 (PF18877), CshA_repeat (PF19076), and Big_13 (PF19077), among others. Phylogenetic trees of bacterial genomes were generated by first selecting all pairs of homologous genes for each genome pair using BLASTn, then computing a genomic sequence identity matrix and finally creating a dendrogram of strains by hierarchical clustering.

## General

We acknowledge Johan Turkenburg and Sam Hart for assistance with crystal testing and data collection, Judith Hawkhead for protein expression and purification, and Andrew Leech, Rachael Cooper and Michael Hodgkinson for *T*_m_ measurements and figure preparation. The authors also thank the Diamond Light Source for access to beamline I03 (proposal number mx-9948) that contributed to the results presented here. This work used the Bioscience Technology Facility at the University of York.

## Author Contributions

J.R.P. proposed the study; F.W., A.L., J.G., S.C.G., C.G.B., E.P., A.N.StJ., H.T.J., A.B. and C.D. performed experiments and analysed data; J.R.P., A.B., C.G.B., M.J.P., J.W.B.M. and E.P. guided experimental design and data analysis. All authors contributed to the interpretation of the data and to the writing of the manuscript.

## Data and materials availability

Atomic models of ΔN-Sgo_R2, Sgo_R10 and Sgo_R3-4 have been deposited in the Protein Data Bank with PDB IDs 7AVJ, 7AVK and 7AVH, respectively. All data needed to evaluate the conclusions in the paper are present in the paper and/or the Supplementary Materials.

