## Supplementary for "Periscope Proteins are variable length regulators of bacterial cell surface interactions"

Supplementary Materials

Figs. S1 to S6
Tables S1 to S3

Legend for Supplementary Data 1

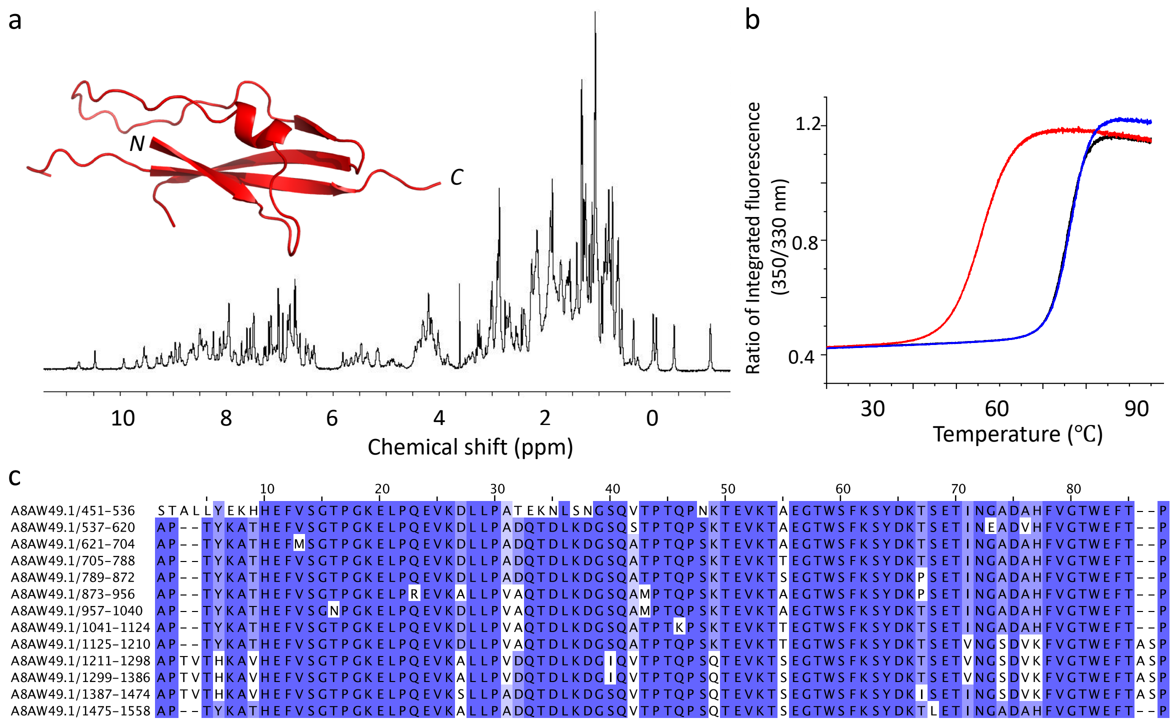


**Figure S1. Defining the structural repeats of Sgo_0707 from *Streptococcus gordonii.***

**(a)** 1D $^{1}$H NMR spectrum of ΔN-Sgo_R2, structure of ΔN-Sgo_R2 inset. **(b)** $T_m$ values for ΔN-Sgo_R2 (red), Sgo_R3 (black) and Sgo_R10 (blue) determined using nanoDSF. **(c)** sequence alignment of the 13 adjacent SHIRT domain repeats in Sgo_0707 protein.

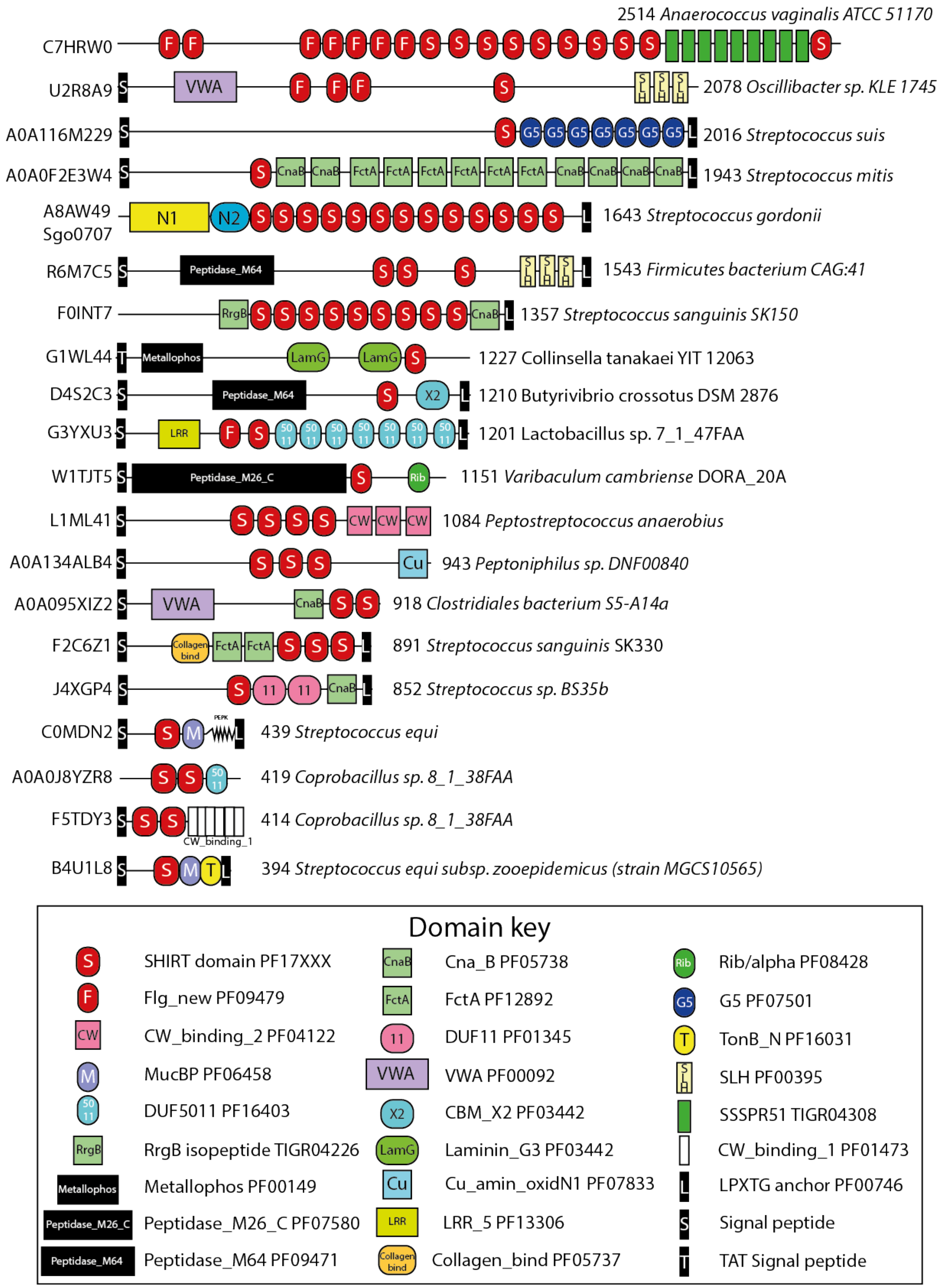


**Figure S2. SHIRT domains are found in many other proteins, often in tandem array.** Each line in the figure shows a representative protein domain architecture containing SHIRT/S domains with UniProt accession number shown at the left. The domain architecture shows boxes and shapes (Domain key) representing various domains, repeats and motifs identified in the proteins using Pfam and TIGRfams. The N1 and N2 domains from Sgo_0707 are included based on the known structure but are not represented within any domain database. Potential enzymatic domains are shown as the larger black boxes with white text, while the localization motifs are shown as small block boxes with a white letter to represent their type. At the end of each line the length of the protein is shown followed by the species and strain if known.

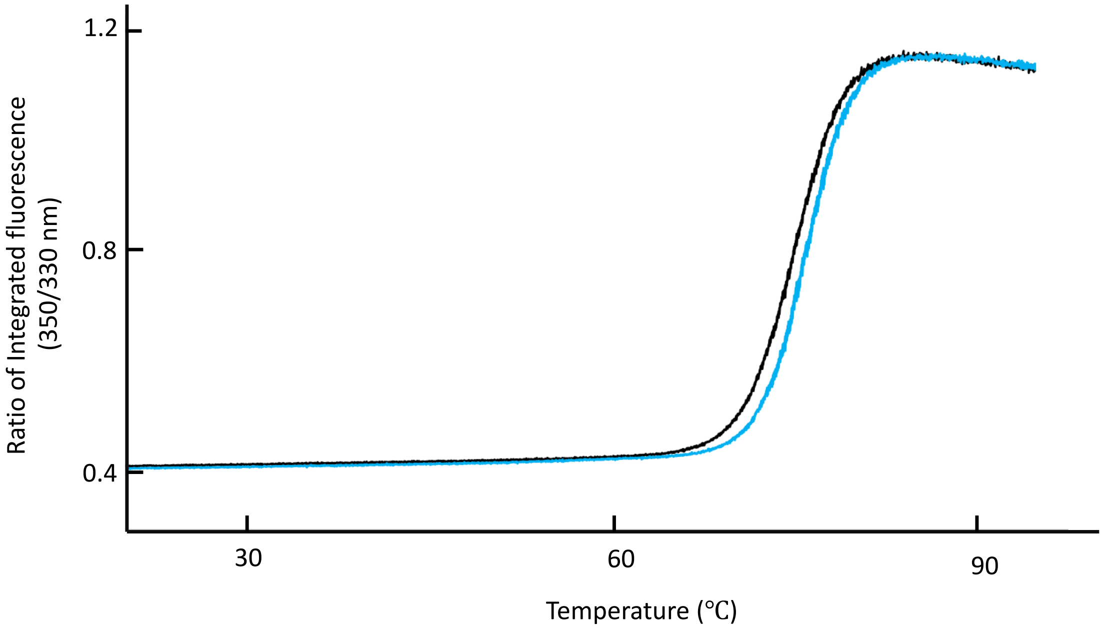


**Figure S3. The interdomain interface in Sgo_R3-4 is very limited.**

$T_m$ analysis using nanoDSF of Sgo_R3 (black; Sgo_R3 in fig. S1C) compared with Sgo_R3-4 (cyan).

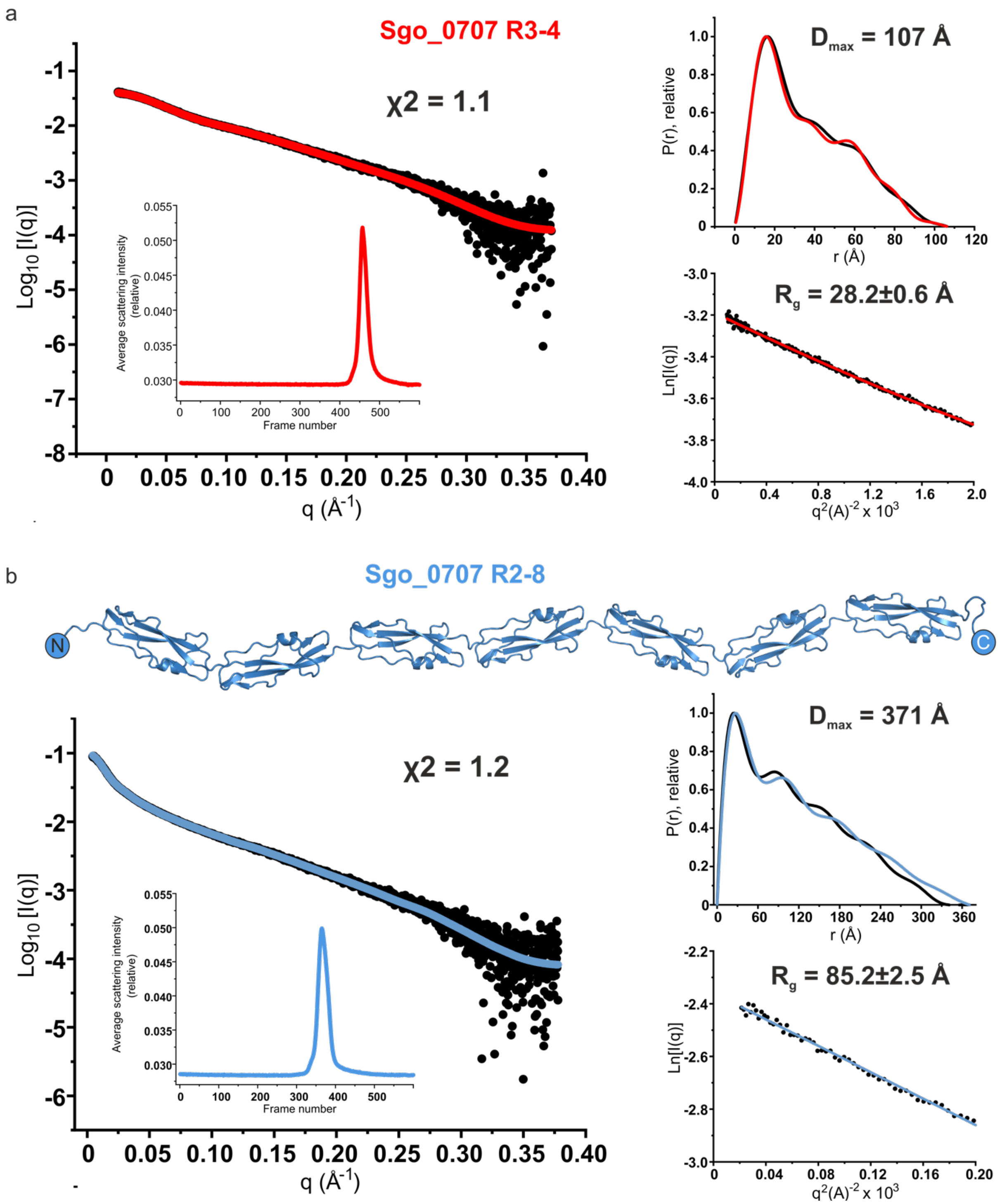


**Figure S4. Validation of Sgo_0707 R3-4 and Sgo_0707 R2-8 models in solution via SAXS.** **(a)** (Left) Fit of Sgo_0707 R3-4 crystal structure (see Fig. 2A) to experimental scattering data. Experimental (black) and model (red) scattering curves are displayed to a maximum scattering angle of $q$ = 0.378 $Å^{-1}$. Scattering data are accounted for by a single Sgo_0707 R3-4 model, with fit value ($\chi$2) displayed. A trace of the in-line SEC run is displayed (inset). Normalised pair-distance distribution function (experimental – black, model – red) is displayed, with the derived maximum intra-particle distance ($D_{max}$) displayed (real space $R_g$ = 30.1±0.1 Å, $I$(0) = 0.04) (top right). The linear portion of the Guinier region confirms monodispersity ($q*R_g$ range = 0.4–1.2; $I(0)$ = 0.04) (bottom right). **(b)** Fit of Sgo_0707 R2-8 model to experimental scattering data. A model for Sgo_R2-8 was generated using iterative placement of the Sgo_R3-4 crystal structure (top; see methods). Experimental (black) and model (blue) scattering curves are displayed to a maximum scattering angle of $q$ = 0.368 Å$^{-1}$ (left). Scattering data are accounted for by a single Sgo_R2-8 model, with fit value displayed. A trace of the in-line SEC run is displayed (inset). Normalised pair-distance distribution function (experimental – black, model – blue) is displayed, with the calculated $D_{max}$ displayed (real space $R_g$ = 96.2±0.5 Å, $I(0)$ = 0.10) (middle right). The Guinier region confirms monodispersity ($q*R_g$ range = 0.5–1.3; $I(0)$ = 0.093) (bottom right).

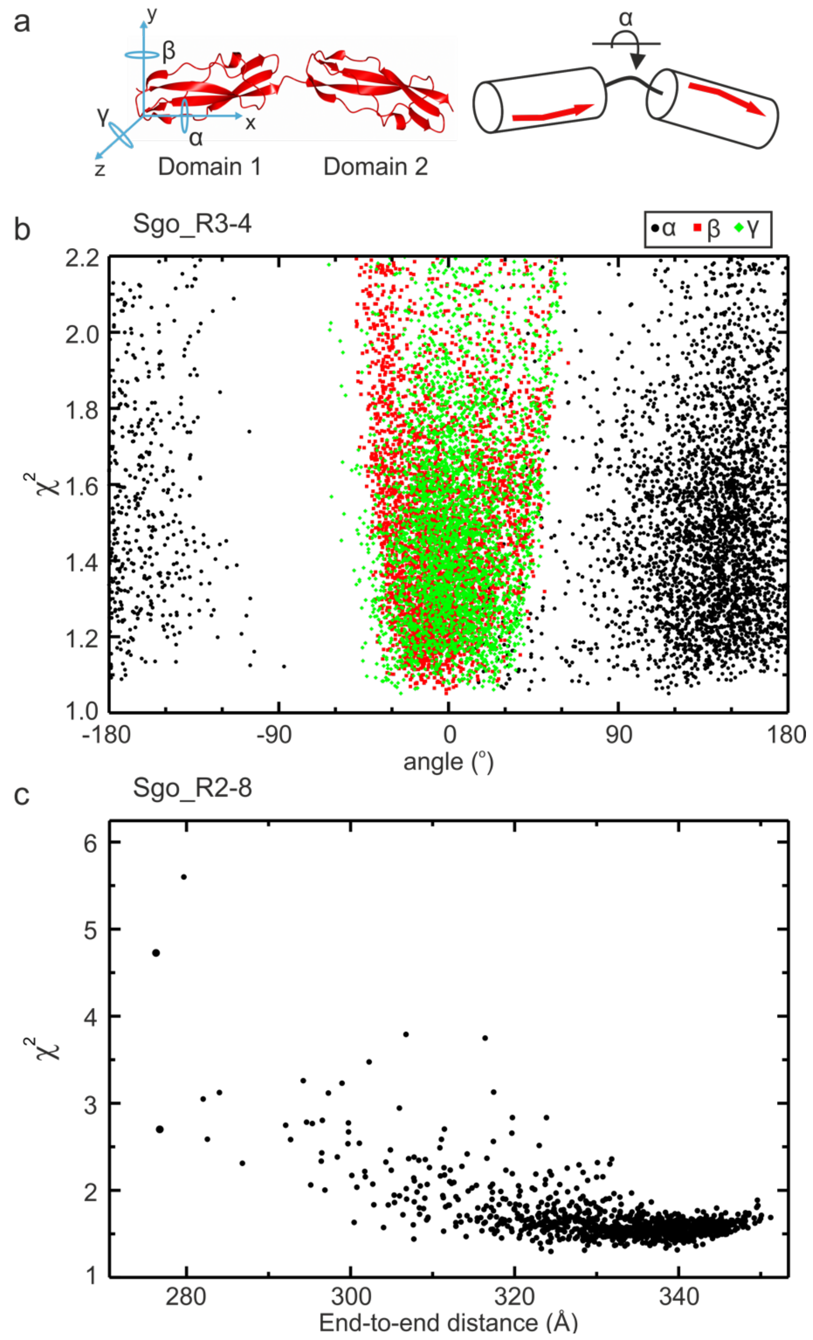


**Figure S5. Elongation of Sgo_0707 is dependent on rotation angle α.**

**(a)** Definition of the three rotation angles (α, β, γ) between consecutive Sgo_0707 domains, along the axes x-, y- and z- (left). Domains can rotate around angle α to maintain elongation (right). **(b)** Fit of simulated Sgo_R3-4 models with variation of all three angles α, β, γ to experimental SAXS data. **(c)** Fit of simulated Sgo_R2-8 models to experimental SAXS data. End-to-end distance is defined as in Fig. 2.

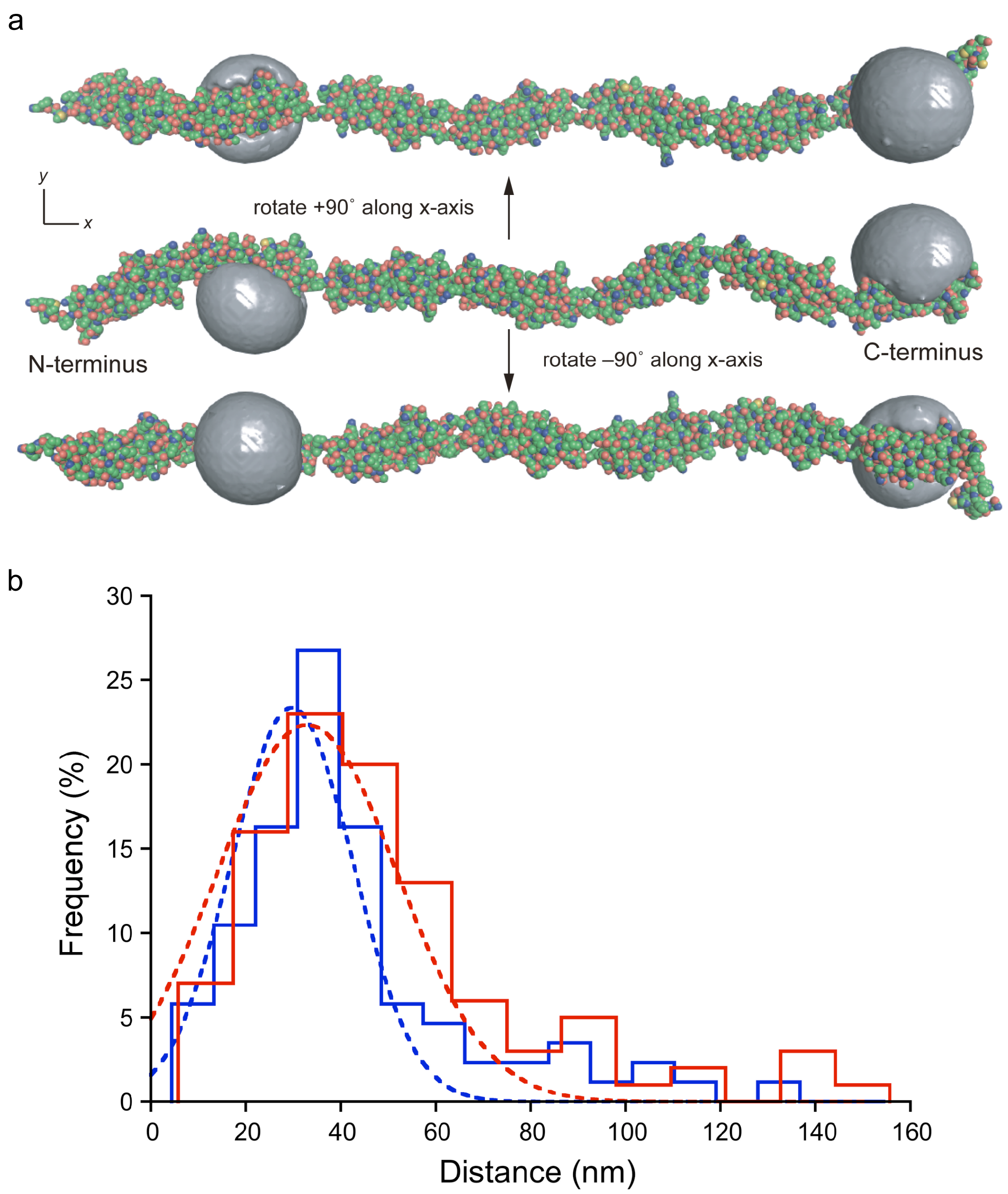


**Figure S6. Determination of inter-dye distances using SHRImP-TIRFM.**

**(a)** The structural schematics show the accessible volume (AV) for both Alexa Fluor 488 (AF488) dyes attached to Sgo_R2-8$^{S666C/S1086C}$ via a $C_5$ linker at positions S666 and S1086. The N- and C-terminal ends of the Sgo_R2-8$^{S666C/S1086C}$ construct are labelled. The SAXS-derived structural model (fig. S4B) was utilised to simulate the AVs (grey envelopes) as described in the Methods section using three radii to estimate the volume of the AF488 dye. In this structural model, the distance between the dye attachment points ($C_\beta$ atoms of S666 and S1086 residues) is 24.0 nm, and the predicted distance between the mean dye positions is 24.1 nm. Images were prepared using the PyMOL Molecular Graphics System (version 2.4.0, Schrödinger LLC). **(b)** SHRImP-TIRFM determination of the inter-dye distances for AF488-labelled Sgo_R2-8$^{S666C/S1086C}$ immobilised on 2 µg/mL (blue) or 20 µg/mL (red) poly-D-lysine treated quartz surfaces in the presence of 100-fold molar excess of unlabelled blocking Sgo_R2-8 protein. Colour-coded, dashed lines indicate Gaussian fits to the histograms for the 2 µg/mL ($n$ = 86, $R^2$ = 0.92) and 20 µg/mL ($n$ = 100, $R^2$ = 0.94) poly-D-lysine-treated quartz surfaces (mean ± standard error = 29.8 ± 1.38 nm and 33.0 ± 1.90 nm, respectively).

*Supplementary Table 1: Data collection and refinement statistics*

| **Data Collection Statistics** | **ΔN-Sgo_R2** | **Sgo_R10** | **Sgo_R3-4** |
|---|---|---|---|
| Wavelength (Å) | 0.85 | 0.78 | 0.976 |
| Space group | $P2_12_12_1$ | $P2_12_12$ | $P2_1$ |
| Cell dimensions<br>a, b, c (Å)<br>$\beta$ (□) | <br>21.4, 40.8, 82.5<br>90.0 | <br>65.4 48.0 48.4<br>90.0 | <br>24.1, 37.2, 101.1<br>95.8 |
| Resolution limits (Å)[1] | 82.5—0.95 (0.96—0.95) | 38.9—0.82<br>(0.85 —0.82) | 37.2—7.39<br>(1.37—1.35) |
| No. reflections<br>Total[1]<br>Unique[1] | <br>281952 (12914)<br>44546 (2604) | <br>292386 (24481)<br>147184 (11281) | <br>157956 (7439)<br>39416 (1911) |
| $R_{merge}$[1,2] | 0.034 (0.681) | 0.045 (2.057) | 0.066 (0.960) |
| Mean $I/\sigma I$[1] | 24.8 (2.3) | 28.3 (0.3) | 8.9 (1.4) |
| Half-set correlation CC (1/2)[1] | 0.999 (0.839) | 1.000 (0.239) | 0.998 (0.562) |
| Wilson B-factor (Å$^2$) | 8.9 | 10.8 | 10.8 |
| Completeness (%)[1] | 95.2 (75.8) | 96.8(76.5) | 99.9 (99.6) |
| Redundancy[1] | 6.3 (5.0) | 2.0(1.9) | 4.0 (3.9) |
| **Refinement Statistics** | | | |
| Resolution (Å)[1] | 43.2 (0.95) | 38.9 (0.82) | 34.9 (1.35) |
| No. of reflections<br>Working<br>Free | <br>46601<br>2309 | <br>144297<br>1454 | <br>37511<br>1892 |
| $R_{work}/R_{free}$ (%) | 12.5/13.3 | 14.7/16.2 | 13.4/17.0 |
| Rms from ideality<br>Bond length (Å)<br>Angles (°) | <br>0.017<br>1.96 | <br>0.017<br>1.48 | <br>0.023<br>2.15 |
| No. of atoms<br>Protein<br>Ligand/ion<br>Water | <br>661<br>-<br>104 | <br>1442<br>15<br>247 | <br>1404<br>33<br>200 |
| Average B (Å$^2$)<br>Protein<br>Ligand/ion<br>Water | <br>14.9<br>-<br>28.3 | <br>18.1<br>19.7<br>27.8 | <br>18<br>22.4<br>34.0 |
| Ramachandran angles (%)<br>Favoured<br>Allowed<br>Outliers | <br>98.0<br>2.0<br>0.0 | <br>98.15<br>1.85<br>0.0 | <br>97.0<br>3.0<br>0.0 |

[1] Values in parentheses are for the highest resolution shell.

[2] $R_{merge} = \Sigma|I - \langle I \rangle| / \Sigma I$

*Supplementary Table 2: Primer sequences for SHIRT domain construct cloning and mutagenesis*

| **Construct** | **Primers** |
|---|---|
| ΔN-Sgo_R2 (aa 544–627) | 5'-TCCAGGGACCAGCAA**TG**CATGAATTTGTGAGCGGCAC-3' and 5'-TGAGGAGAAGGCGCGTTAGGTGGCTTTGTAGGTCGG-3' |
| Sgo_R3 (aa 621–705) | 5'-TCCAGGGACCAGCA**ATG**GCCCCGACCTACAAAGCCACC-3' and 5'-TGAGGAGAAGGCGCGTTAGGCCGGGGTAAATTCCCAGGTGC-3' |
| Sgo_R10 (aa 1211–1299) | 5'-TCCAGGGACCAGCA**ATG**GCTCCAACAGTGACTC-3' and 5'-TGAGGAGAAGGCGCGTTATTAAGCTGGGCTCGCTG-3' |
| Sgo_R3-4 (aa 621–789) | TCCAGGGACCAGCA**ATG**GCCCCGACCTACAAAGCCACC-3' and 5'-TGAGGAGAAGGCGCGTTATGCCGGGGTGAATTCCCAGGTAC-3' |
| Sgo_R2-8 S666C | 5'-GACTCAACCTTGTAAAACAG-3' and 5'-CTGTTTTACAAGGTTGAGTC-3' |
| Sgo_R2-8 S1086C | 5'-GACAAAACCCTGTAAGACCG-3' and 5'-CGGTCTTACAGGGTTTTGTC-3' |

*Supplementary Table 3: Synthetic DNA sequences used in this study*

| | |
|---|---|
| Sgo_0707 residues 544–795 | catgaatttgtgagcggcaccccgggcaaggaactgcctcaggaagtgaaggacctgctgccg<br>gccgatcaaaccgatctgaaagacggcagtcagagcaccccgacccagccgagtaaaaccgag<br>gtgaaaaccgccgaaggcacctggagctttaagagctacgacaagaccagcgaaaccatcaac<br>gaagccgatgtgcactttgtgggtacctgggagttcaccccggccccgacctacaaagccacc<br>catgagttcatgagcggcacaccgggtaaagaactgcctcaagaggtgaaggatctgctgcct<br>gcagatcaaaccgacctgaaggatggcagtcaagccaccccgacacagccgagcaaaacagaa<br>gtgaagacagccgagggcacctggagcttcaaaagctatgacaaaaccagcgagaccattaac<br>ggcgcagatgcccactttgtgggcacctgggaatttaccccggccccgacatacaaggccacc<br>cacgagtttgtgagcggtacaccgggcaaagaactgccgcaggaagttaaagatctgctgccg<br>gccgaccagaccgacctgaaagatggtagccaggcaaccccgacccaaccgagcaagacagaa<br>gtgaaaaccaccgaaggcacctggagcttcaagagttatgataagaccagcgaaaccattaat<br>ggcgccgatgcccattttgttggtacctgggaattcaccccggcaccgacctataaggccacc |
| Sgo_R2-8 | gcgccaacctacaaggcaactcacgagtttgtcagcggaactccaggaaaagaacttcca<br>caagaagtgaaggacctgcttccagcagaccaaacagacttgaaagatggtagtcaatcg<br>actccaacgcaaccaagtaaaaccgaggttaagacagcagaaggcacatggagcttcaag<br>tcctatgacaagacttccgaaaccatcaatgaagcagacgtacacttcgtaggaacatgg<br>gaattcaccccagcgccaacctacaaggcgactcatgagtttatgagtggaaccccaggt<br>aaagagcttccacaagaagtgaaagacctgcttccagcagaccaaacagacttgaaagat<br>ggaagccaagcgactccaactcaaccaagtaaaacggaagttaagacagcagaaggcact<br>tggagtttcaagtcatacgacaagacttctgaaaccattaatggcgcggacgcacacttc<br>gtaggcacatgggaattcaccccagcgccaacctacaaggcgactcatgagtttgtgagt<br>ggaaccccaggtaaagagcttccacaagaagtgaaagacctgcttccagcagaccaaaca<br>gacttgaaagatggaagtcaagccactccaacgcaaccaagtaaaacggaagtgaagacg<br>acagaaggtacttggagtttcaagtcatacgacaagacttctgaaaccattaatggcgcg<br>gacgcacacttcgtaggcacatgggaattcaccccagcgccaacctacaaggcgactcat<br>gagtttgtgagtggaaccccaggtaaagagcttccacaagaagtgaaagacctgcttcca<br>gcagaccaaacagacttgaaagatggaagccaagcgactccaacacaaccaagtaaaacg<br>gaagttaagacgtcagaaggcacttggagcttcaagtcctatgacaagccgtctgaaacc<br>atcaatggagcagacgcacacttcgtaggcacctgggaattcaccccagcgccaacctac<br>aaggcgacacacgagtttgtcagcggaactccaggcaaagagcttccacgagaagtaaaa<br>gcactgcttccagtagctcaaacagacttgaaagatggtagccaagcgatgccaacgcaa<br>ccaagtaaaacagaggttaagacagcagaaggcacttggagcttcaagtcctatgacaag<br>ccgtctgaaaccatcaatggagcagacgcacactttgtcggtacttgggaatttacccca<br>gcaccaacctacaaggcaacacacgagtttgtgagtggaaatccaggtaaagagcttcca<br>caagaagtaaaagatctgcttccagtagctcaaacagacttgaaagatggtagtcaagcg<br>atgccaacgcaaccaagtaaaacggaagttaagacagcagaaggtacttggagtttcaaa<br>tcatacgataagacttccgaaaccatcaatggagcagacgcacactttgtaggaacatgg<br>gaattcaccccagcgccaacctacaaggcgacacacgagtttgtcagcggaactccaggc<br>aaagagcttccacaagaagtaaaagatctgcttccagtagctcaaacagacttgaaagat<br>ggtagccaagcaacaccaacaaaaccaagtaaaacggaagtgaagacaactgaaggcact<br>tggagtttcaaatcatacgataagacttccgaaaccatcaatggagcagacgcacacttt<br>gtaggaacatgggaattcaccccagcg |
| Sgo_R10 | gctccaacagtgactcataaagcagttcacgaatttgtgagtggaactccaggcaaagag<br>cttccacaagaagtgaaagccctgcttccagtagatcaaacagatctgaaagatggcatt<br>caagtgactccaacacaaccaagtcaaacagaggttaagacatcagaaggcacatggagc<br>ttcaagtcatacgataagacttcagagactgtcaacggttcagatgttaagtttgtagga<br>acatgggaatttacagcgagcccagcttaa |

**Supplementary Data 1** Table of Periscope proteins identified from the NCTC3000 dataset. Columns names stand for: cluster identifiers (clustid), recognisable name from gene names and other literature references or UNK for unknown name (name), number of proteins in the cluster (size), minimum number of repeats (minrep), maximum number of repeats (maxrep), length of the repeat (replen), average percentage of protein sequence identity of repeats (psim), average percentage of DNA sequence identity of repeats (psim.dna), closest UniProt protein (uniprotid), Pfam families in the protein (pfam), Pfam families of the repeating sequence (pfam.rep), NCTC3000 species where the protein was found (species), and GO terms for location (GO_ location), function (GO_function) and process (GO_process).

| clustid | name | uniprotid | size | replen | pfam.rep | minrep | maxrep | psim | psim.dna | pfam | species | GO_location | GO_function |
|---|---|---|---|---|---|---|---|---|---|---|---|---|---|
| 1 | Spa | M4MB15 | 97 | 59 | B | 2 | 7 | 0.86541237 | 0.84891304 | B, Gram_pos_anchor, LysM, Octapeptide, YSIRK_signal | Staphylococcus argenteus, Staphylococcus aureus, Staphylococcus aureus MRSA 252, Staphylococcus sp., Staplylococcus schweitzeri | cell wall [GO:0005618]; extracellular region [GO:0005576]; membrane [GO:0016020] | immunoglobulin binding [GO:0019865] |
| 2 | UNK_DUF1542 | | 87 | 78 | DUF1542 | 2 | 4 | 0.94965517 | 0.94264368 | DUF1542, FIVAR, GA, Rib, YSIRK_signal | Staphylococcus aureus, Staphylococcus aureus MRSA 252, Staphylococcus sp., Staplylococcus schweitzeri | | |
| 3 | | A0A0H2WZC2 | 67 | 53 | | 2 | 4 | 0.87731343 | 0.94776119 | | Staphylococcus aureus, Staphylococcus sp. | integral component of membrane [GO:0016021] | |
| 4 | SasG | Q2G2B2 | 34 | 128 | G5, SasG_E | 3 | 13 | 0.84470588 | 0.97764706 | Bact_lectin, G5, Gram_pos_anchor, SasG_E, YSIRK_signal | Staphylococcus aureus | cell wall [GO:0005618]; extracellular region [GO:0005576]; membrane [GO:0016020] | |
| 5 | CdrA | A0A485CMM4 | 24 | 81 | MBG_2 | 3 | 15 | 0.88229167 | 0.86541667 | Haemagg_act, MBG_2 | Pseudomonas aeruginosa | | |
| 6 | BapA | A0A5E9KBQ0 | 23 | 83 | Big_6 | 2 | 15 | 0.89268116 | 0.82724638 | Big_6, FG-GAP | Pseudomonas aeruginosa | | |
| 7 | UNK_Big13 | Q0B1Q8 | 17 | 103 | Big_13 | 2 | 31 | 0.93392157 | 0.98588235 | Big_9, Big_13, Cadherin, Cadherin_3, Cadherin_4, HemolysinCabind, VWA_2 | Vibrio vulnificus | | sialate O-acetylesterase activity [GO:0001681] |
| 8 | ShdA | A0A3U7M2R1 | 15 | 59 | PATR | 4 | 6 | 0.85066667 | 0.902 | Autotransporter, ESPR, PATR | Salmonella, Salmonella enterica, Salmonella enterica subsp. enterica, Salmonella enterica subsp. enterica serovar Java, Salmonella enterica subsp. enterica serovar Typhimurium, Salmonella enteritidis, Salmonella sp. | integral component of membrane [GO:0016021]; outer membrane [GO:0019867] | |
| 9 | SasY | A0A2T4PXZ6 | 14 | 87 | Big_6 | 2 | 21 | 0.92285714 | 0.94142857 | Big_6, Gram_pos_anchor, YSIRK_signal | Staphylococcus aureus, Staphylococcus epidermidis, Staphylococcus warneri | cell wall [GO:0005618]; extracellular region [GO:0005576] | |
| 10 | SraP | A0A376H473 | 14 | 98 | | 2 | 46 | 0.91761905 | 0.99285714 | Bact_lectin, Gram_pos_anchor, YSIRK_signal | Streptococcus agalactiae, Streptococcus canis | cell wall [GO:0005618]; extracellular region [GO:0005576]; integral component of membrane [GO:0016021] | |
| 11 | | A0A3X6MKL6 | 13 | 56 | Phage_fiber_2 | 2 | 4 | 0.84615385 | 0.86 | DUF3751, Phage_fiber_2 | Salmonella, Salmonella enterica, Salmonella enterica subsp. enterica, Salmonella enterica subsp. enterica serovar Goldcoast, Salmonella enterica subsp. enterica serovar Typhimurium, Salmonella enterica subsp. salamae, Salmonella sp | | |
| 12 | Esp | Q9Z4N7 | 12 | 82 | Rib | 3 | 10 | 0.87208333 | 0.96916667 | Big_6, Gram_pos_anchor, Rib, YSIRK_signal | Enterococcus faecalis, Enterococcus faecium, Enterococcus hirae, Streptococcus | cell wall [GO:0005618]; extracellular region [GO:0005576]; membrane [GO:0016020] | |
| 13 | UNK_YadA | A0A3T3ETZ9 | 12 | 90 | YadA_stalk | 3 | 4 | 0.85916667 | 0.9125 | ESPR, YadA_anchor, YadA_head, YadA_stalk | Salmonella enterica, Salmonella enterica subsp. enterica, Salmonella enterica subsp. enterica serovar Madelia, Salmonella enterica subsp. enterica serovar Oranienburg | cell outer membrane [GO:0009279]; cell surface [GO:0009986]; integral component of membrane [GO:0016021] | |
| 14 | | A0A449F5R6 | 11 | 53 | | 2 | 6 | 0.89272727 | 0.952 | | Streptococcus agalactiae | | |
| 15 | | P37710 | 11 | 68 | LysM | 4 | 6 | 0.86636364 | 0.87272727 | Glucosaminidase, LysM | Enterococcus faecalis, Streptococcus | | |
| 16 | | A0A486RRH2 | 10 | 79 | | 3 | 4 | 0.909 | 0.97 | | Klebsiella pneumoniae, Klebsiella sp, Klebsiella sp. | | |
| 17 | ZmpC | F0I817 | 10 | 96 | G5 | 2 | 10 | 0.892 | 0.82666667 | FIVAR, G5, Peptidase_M26_C, Peptidase_M26_N, YSIRK_signal | Streptococcus gordonii, Streptococcus sanguinis, Streptococcus sp. | cell wall [GO:0005618]; extracellular region [GO:0005576]; integral component of membrane [GO:0016021] | metalloendopeptidase activity [GO:0004222]; zinc ion binding [GO:0008270] |
| 18 | Epf | Q48UF3 | 10 | 81 | DUF1542 | 3 | 10 | 0.8885 | 0.9145 | DUF1542, Gram_pos_anchor | Streptococcus pyogenes | cell wall [GO:0005618]; extracellular region [GO:0005576]; integral component of membrane [GO:0016021] | |
| 19 | Spg | A0A4V0F6C4 | 10 | 75 | GA-like | 2 | 4 | 0.845 | 0.98 | GA-like, Gram_pos_anchor, IgG_binding_B | Streptococcus dys, Streptococcus dysgalactiae subsp. equisimilis, Streptococcus sp. | cell wall [GO:0005618]; extracellular region [GO:0005576]; integral component of membrane [GO:0016021] | |
| 20 | CshA | A0A0F2D244 | 10 | 101 | CshA_repeat | 3 | 14 | 0.839 | 0.822 | CshA_NR2, CshA_repeat, Gram_pos_anchor, YSIRK_signal | Streptococcus gordonii, Streptococcus sp. | cell wall [GO:0005618]; extracellular region [GO:0005576]; membrane [GO:0016020] | |
| 21 | UNK | A0A507HZU9 | 9 | 154 | | 3 | 4 | 0.91444444 | 0.9 | Gram_pos_anchor | Enterococcus faecalis, Streptococcus | | |
| 22 | UNK_Lyase | A0A449G498 | 8 | 70 | Lyase_8_C | 3 | 7 | 0.89125 | 0.94625 | Big_2, F5_F8_type_C, FIVAR, Gram_pos_anchor, Lyase_8, Lyase_8_C, Lyase_8_N | Enterococcus faecalis, Streptococcus | | |
| 23 | PavB_2 | A0A3R9HDH8 | 8 | 150 | SSURE | 2 | 6 | 0.8875 | 0.90285714 | Gram_pos_anchor, SSURE, YSIRK_signal | Streptococcus cristatus, Streptococcus gordonii, Streptococcus sanguinis, Streptococcus sp. | cell wall [GO:0005618]; extracellular region [GO:0005576]; integral component of membrane [GO:0016021] | |
| 24 | ShdA_2 | A0A400S776 | 8 | 64 | PATR | 2 | 14 | 0.88625 | 0.85261905 | ESPR, PATR | Salmonella enterica, Salmonella enterica subsp. enterica, Salmonella enterica subsp. enterica serovar Goldcoast | integral component of membrane [GO:0016021]; outer membrane [GO:0019867] | |
| 25 | R28 | Q48S64 | 7 | 79 | Rib | 2 | 15 | 0.95571429 | 0.99428571 | AlphaC_C, AlphaC_N, Gram_pos_anchor, Rib, YSIRK_signal | Streptococcus agalactiae, Streptococcus pyogenes, Streptococcus sp. | cell wall [GO:0005618]; extracellular region [GO:0005576]; membrane [GO:0016020] | |
| 26 | Sgo0707 | A0A3R9JLL4 | 7 | 84 | SHIRT | 7 | 13 | 0.91428571 | 0.89571429 | Gram_pos_anchor, SHIRT | Streptococcus gordonii, Streptococcus sp. | integral component of membrane [GO:0016021] | |
| 27 | UNK_SpaA | A0A455TW44 | 7 | 93 | SpaA | 2 | 4 | 0.89285714 | 0.95142857 | SpaA | Enterococcus faecium, Enterococcus hirae, Streptococcus sp. | integral component of membrane [GO:0016021] | |
| 28 | UNK_SpaA | A0A2J9ETF2 | 7 | 112 | SpaA | 3 | 17 | 0.87142857 | 0.90428571 | Big_8, Gram_pos_anchor, SpaA | Streptococcus gordonii, Streptococcus sp. | cell wall [GO:0005618]; extracellular region [GO:0005576] | |
| 29 | UNK | A0A381IWY6 | 6 | 96 | | 3 | 9 | 0.93833333 | 0.94333333 | Gram_pos_anchor | Clostridium perfringens | cell wall [GO:0005618]; extracellular region [GO:0005576] | |
| 30 | | A0A2K9C0W2 | 6 | 101 | | 3 | 21 | 0.91166667 | 0.99166667 | Cadherin_3, HemolysinCabind, Peptidase_M10_C, VWA | Vibrio mimicus | | calcium ion binding [GO:0005509] |
| 31 | SasY_2 | A0A4V0BAN3 | 6 | 87 | Big_6 | 3 | 15 | 0.90833333 | 0.99666667 | Big_6, YSIRK_signal | Staphylococcus aureus, Staphylococcus sp. | membrane [GO:0016020] | |
| 32 | UNK_G5 | A0A4U9ZZC2 | 6 | 79 | G5 | 3 | 5 | 0.90333333 | 0.89833333 | G5, Gram_pos_anchor | Streptococcus gordonii, Streptococcus sp. | cell wall [GO:0005618]; extracellular region [GO:0005576]; integral component of membrane [GO:0016021] | |
| 33 | UNK_Big13 | A0A2I5HP40 | 6 | 103 | Big_13 | 2 | 9 | 0.89333333 | 0.97666667 | Big_3_2, Big_3_3, Big_6, Big_13 | Salmonella enterica subsp. arizonae, Salmonella enterica subsp. houtenae | | hydrolase activity [GO:0016787] |
| 34 | | A0A2X3SFE2 | 6 | 51 | | 3 | 5 | 0.885 | 0.935 | | Streptococcus equi, Streptococcus equi subsp zooepidemicus, Streptococcus equi subsp. zooepidemicus, Streptococcus sp, Streptococcus sp. | | |
| 35 | | P71397 | 6 | 56 | | 2 | 6 | 0.87833333 | 0.91666667 | Complex1_51K, Fer4_10, Fer4_17, Fer4_7, Fer4_8, Fer4_9, NQRA, RnfC_N, SLBB | Haemophilus influenzae | plasma membrane [GO:0005886] | 4 iron, 4 sulfur cluster binding [GO:0051539]; electron transfer activity [GO:0009055]; metal ion binding [GO:0046872] |
| 36 | | A0A2L2PB53 | 6 | 103 | GBS_Bsp-like | 3 | 4 | 0.855 | 0.93666667 | CHAP, GBS_Bsp-like, Glyco_hydro_25 | Streptococcus gordonii, Streptococcus sp. | | lysozyme activity [GO:0003796] |
| 37 | SasG_2 | T1YDP9 | 5 | 128 | G5, SasG_E | 3 | 13 | 0.936 | 0.978 | Bact_lectin, G5, Gram_pos_anchor, SasG_E, YSIRK_signal | Staphylococcus argenteus, Staphylococcus aureus, Staplylococcus schweitzeri | cell wall [GO:0005618]; extracellular region [GO:0005576]; membrane [GO:0016020] | beta-N-acetylhexosaminidase activity [GO:0004563]; N-acetyl-beta-D-galactosaminidase activity [GO:0102148] |
| 38 | Fap | A0A1Q1FVK8 | 5 | 131 | DUF11 | 2 | 5 | 0.932 | 0.938 | DUF11, Gram_pos_anchor | Enterococcus faecalis | cell wall [GO:0005618]; extracellular region [GO:0005576]; integral component of membrane [GO:0016021] | |
| 39 | Rib | A0A1A9DZG2 | 5 | 82 | Rib | 2 | 7 | 0.916 | 0.944 | AlphaC_C, AlphaC_N, Gram_pos_anchor, Rib, YSIRK_signal | Streptococcus agalactiae | cell wall [GO:0005618]; extracellular region [GO:0005576]; membrane [GO:0016020] | |
| 40 | PavB | C1C9Y2 | 5 | 152 | SSURE | 2 | 9 | 0.912 | 0.986 | Gram_pos_anchor, SSURE, YSIRK_signal | Streptococcus pneumoniae | cell wall [GO:0005618]; extracellular region [GO:0005576]; integral component of membrane [GO:0016021] | |
| 41 | | A0A4V0A1I1 | 5 | 59 | | 2 | 4 | 0.874 | 0.85666667 | | Streptococcus equisimilis, Streptococcus sp. | | |
| 42 | | A0A3Z9J190 | 5 | 56 | Phage_fiber_2 | 4 | 5 | 0.868 | 0.932 | DUF3751, Phage_fiber_2, PTR | Escherichia coli | | |
| 43 | UNK_MucBP | A0A5S4TIX9 | 5 | 81 | MucBP | 4 | 7 | 0.853 | 0.893 | Big_8, Fn_bind, Gram_pos_anchor, MucBP | Streptococcus dys, Streptococcus equisimilis, Streptococcus sp. | cell wall [GO:0005618]; extracellular region [GO:0005576]; integral component of membrane [GO:0016021] | |
| 44 | UNK_Invasin | A0A376HGL9 | 5 | 97 | Big_1 | 3 | 12 | 0.84333333 | 0.856 | Big_1, IAT_beta, Invasin_D3 | Escherichia coli | | |
| 45 | UNK_Big3 | E2Z005 | 5 | 73 | Big_3, DUF5011 | 3 | 6 | 0.826 | 0.782 | Big_3, DUF5011, SpaA | Enterococcus faecalis, Streptococcus | integral component of membrane [GO:0016021] | |
| 46 | Bca_2 | A0A4U9ZPF1 | 4 | 134 | Rib | 9 | 24 | 0.975 | 0.9925 | Flg_new, Gram_pos_anchor, Rib, YSIRK_signal | Streptococcus sp. | cell wall [GO:0005618]; extracellular region [GO:0005576]; integral component of membrane [GO:0016021] | calcium ion binding [GO:0005509] |
| 47 | | A0A376Y364 | 4 | 99 | | 19 | 43 | 0.9675 | 0.9975 | | Escherichia coli | | |
| 48 | | A0A426MI08 | 4 | 87 | | 2 | 3 | 0.9275 | 0.9525 | PhageMin_Tail | Escherichia coli | integral component of membrane [GO:0016021] | |
| 49 | UNK_Big6 | A0A0M2AG42 | 4 | 90 | Big_6 | 4 | 5 | 0.875 | 0.95 | Big_6, Gram_pos_anchor | Enterococcus faecalis | cell wall [GO:0005618]; extracellular region [GO:0005576]; integral component of membrane [GO:0016021] | |
| 50 | LytD | A0A2X2YDI6 | 4 | 85 | SH3_3 | 3 | 6 | 0.87375 | 0.9525 | Glucosaminidase, SH3_3 | Clostridium perfringens | | amidase activity [GO:0004040]; mannosyl-glycoprotein endo-beta-N-acetylglucosaminidase activity [GO:0033925] |
| 51 | UNK_Big13 | A0A4P9IZ13 | 3 | 95 | Big_13 | 9 | 16 | 0.95 | 0.99666667 | Big_3_2, Big_9, Big_13, Cadherin_4, HemolysinCabind, VWA_2 | Neisseria elongata | | |
| 52 | SraP_2 | A0A380G3G2 | 3 | 89 | He_PIG | 2 | 14 | 0.94333333 | 0.98 | Big_8, Gram_pos_anchor, He_PIG, SdrG_C_C, YSIRK_signal | Staphylococcus saprophyticus, Staphylococcus saprophyticus subsp. saprophyticus | cell wall [GO:0005618]; extracellular region [GO:0005576]; membrane [GO:0016020] | calcium ion binding [GO:0005509] |
| 53 | YeeJ | A0A192CME6 | 3 | 119 | Invasin_D3 | 2 | 3 | 0.93666667 | 0.98 | Big_1, IAT_beta, Invasin_D3 | Escherichia coli | | |
| 54 | UNK_Rib | A0A2T4QWP0 | 3 | 86 | Rib | 5 | 12 | 0.92333333 | 0.98666667 | Big_6, Gram_pos_anchor, Rib, YSIRK_signal | Staphylococcus hominis | cell wall [GO:0005618]; extracellular region [GO:0005576]; integral component of membrane [GO:0016021] | |
| 55 | UNK_Rib | H2A912 | 3 | 79 | Rib | 5 | 14 | 0.90666667 | 0.99333333 | Gram_pos_anchor, He_PIG, Rib, YSIRK_signal | Streptococcus agalactiae, Streptococcus sp. | cell wall [GO:0005618]; extracellular region [GO:0005576]; membrane [GO:0016020] | calcium ion binding [GO:0005509] |
| 56 | | A0A3S7N4X5 | 3 | 67 | | 2 | 3 | 0.87666667 | 0.90333333 | Complex1_51K, Fer4_10, Fer4_4, Fer4_7, Fer4_9, NQRA, RnfC_N, SLBB | Mannheimia haemolytica | plasma membrane [GO:0005886] | 4 iron, 4 sulfur cluster binding [GO:0051539]; electron transfer activity [GO:0009055]; metal ion binding [GO:0046872] |
| 57 | UNK_DUF1542 | A0A5S4TRV8 | 3 | 81 | DUF1542 | 5 | 7 | 0.865 | 0.91333333 | DUF1542, Gram_pos_anchor | Streptococcus pyogenes | cell wall [GO:0005618]; extracellular region [GO:0005576]; integral component of membrane [GO:0016021] | |
| 58 | UNK_Lyase | A0A4U9PE62 | 3 | 69 | FIVAR | 5 | 6 | 0.86333333 | 0.92333333 | Big_2, F5_F8_type_C, FIVAR, Lyase_8, Lyase_8_C, Lyase_8_N | Enterococcus faecium, Enterococcus hirae | | |
| 59 | Emm23 | Q9RHV2 | 3 | 63 | | 2 | 3 | 0.86333333 | 0.99 | Gram_pos_anchor, M, YSIRK_signal | Streptococcus pyogenes, Streptococcus sp. | cell wall [GO:0005618]; extracellular region [GO:0005576]; integral component of membrane [GO:0016021] | |
| 60 | | H7C722 | 3 | 93 | | 5 | 7 | 0.84333333 | 0.99 | | Enterococcus cecorum, Enterococcus faecalis | | |
| 61 | | A0A0K0H9Z7 | 3 | 63 | | 3 | 5 | 0.83 | 0.85 | | Salmonella enterica subsp. arizonae | | |
| 62 | UNK_MucBP | A0A0E2I0J3 | 3 | 75 | MucBP | 2 | 3 | 0.82666667 | 0.88333333 | Gram_pos_anchor, MucBP | Enterococcus faecium, Enterococcus hirae | extracellular region [GO:0005576]; integral component of membrane [GO:0016021] | |
| 63 | Bag | A0A4V0C2I4 | 2 | 89 | CFSR | 11 | 12 | 1 | 0.99 | CFSR, Collagen, Gram_pos_anchor, YSIRK_signal | Streptococcus agalactiae, Streptococcus sp. | extracellular region [GO:0005576]; membrane [GO:0016020] | |
| 64 | | A0A3S4N8L4 | 2 | 98 | | 21 | 29 | 0.975 | 0.99 | Big_9, Cadherin_4, Cadherin_5, HemolysinCabind, VWA, VWA_2 | Shewanella putrefaciens | | calcium ion binding [GO:0005509] |
| 65 | PavB_3 | A0A1X1K6M7 | 2 | 149 | SSURE | 2 | 4 | 0.945 | 0.915 | Gram_pos_anchor, SSURE, YSIRK_signal | Streptococcus sp., Streptococcus viridans | cell wall [GO:0005618]; extracellular region [GO:0005576]; integral component of membrane [GO:0016021] | |
| 66 | Rib_2 | A0A4V0EE13 | 2 | 76 | Rib | 7 | 12 | 0.915 | 0.99 | AlphaC_C, AlphaC_N, Gram_pos_anchor, Rib, YSIRK_signal | Streptococcus canis, Streptococcus sp. | cell wall [GO:0005618]; extracellular region [GO:0005576]; membrane [GO:0016020] | |
| 67 | UNK_DUF1542 | A0A4Q9WP12 | 2 | 77 | DUF1542 | 5 | 6 | 0.905 | 0.865 | DUF1542, Gram_pos_anchor, YSIRK_signal | Staphylococcus hominis, Staphylococcus saprophyticus | cell wall [GO:0005618]; extracellular region [GO:0005576]; membrane [GO:0016020] | |
| 68 | UNK_Invasin | A0A377P116 | 2 | 101 | Invasin_D3 | 3 | 6 | 0.9 | 0.96 | Big_1, IAT_beta, Invasin_D3 | Edwardsiella tarda | | |
| 69 | UNK_SpaA | A0A3R9LMV6 | 2 | 114 | SpaA | 7 | 8 | 0.895 | 0.815 | Big_8, Collagen_bind, SpaA | Streptococcus sanguinis | cell wall [GO:0005618]; extracellular region [GO:0005576] | |
| 70 | | A0A5F2FBF2 | 2 | 59 | | 2 | 3 | 0.895 | 0.96 | | Klebsiella oxytoca, Klebsiella sp | | |
| 71 | | A0A5F0TZB4 | 2 | 105 | | 7 | 12 | 0.88 | 0.83 | | Staphylococcus warneri | | |
| 72 | | A0A0H3MI02 | 2 | 56 | Phage_fiber_2 | 3 | 7 | 0.87 | 0.905 | Collar, Phage_fiber_2 | Escherichia coli | | |
| 73 | Vwbl | Q6QQB0 | 2 | 67 | SSSPR-51 | 7 | 9 | 0.87 | 0.84 | Gram_pos_anchor, MucBP, SSSPR-51, YSIRK_signal | Staphylococcus lugdenensis, Staphylococcus lugdunensis | cell wall [GO:0005618]; extracellular region [GO:0005576]; membrane [GO:0016020] | |
| 74 | | A0A2Z6BTL7 | 2 | 83 | | 2 | 6 | 0.865 | 0.965 | Glyco_hydro_25, GW | Enterococcus faecalis | | lysozyme activity [GO:0003796] |
| 75 | | A0A4U8CJT1 | 2 | 103 | | 2 | 5 | 0.865 | 0.98 | Big_3_2, Big_3_3, Big_6 | Salmonella enterica, Salmonella enterica subsp. enterica | | |
| 76 | | A0A4V4RZJ2 | 2 | 100 | Amidase_3 | 5 | 6 | 0.865 | 0.935 | Amidase_3, GBS_Bsp-like | Streptococcus sp. | | N-acetylmuramoyl-L-alanine amidase activity [GO:0008745] |
| 77 | | D7VRT5 | 2 | 63 | | 3 | 4 | 0.86 | 0.925 | E1-E2_ATPase, Hydrolase, Hydrolase_3 | Sphingobacterium spiritivorum | integral component of membrane [GO:0016021]; plasma membrane [GO:0005886] | ATPase-coupled cation transmembrane transporter activity [GO:0019829]; ATP binding [GO:0005524]; hydrolase activity [GO:0016787]; metal ion binding [GO:0046872] |
| 78 | | A0A2X3YLW4 | 2 | 103 | GBS_Bsp-like | 5 | 7 | 0.855 | 0.795 | CHAP, GBS_Bsp-like, Glyco_hydro_25 | Streptococcus sanguinis | | lysozyme activity [GO:0003796] |
| 79 | | A0A0E2HVS0 | 2 | 67 | | 3 | 4 | 0.85 | 0.92 | | Enterococcus faecium, Enterococcus hirae | | |
| 80 | | A0A378K3C6 | 2 | 165 | | 5 | 6 | 0.85 | 0.85 | HemolysinCabind, VWA_2 | Legionella pneumophila | | calcium ion binding [GO:0005509] |
| 81 | UNK_SHIRT | A0A1X1FZN4 | 2 | 84 | Cna_B | 4 | 7 | 0.84 | 0.805 | Cna_B, SHIRT | Streptococcus sanguinis | integral component of membrane [GO:0016021] | |
| 82 | | A0A380M5K5 | 2 | 103 | GBS_Bsp-like | 4 | 5 | 0.83 | 0.775 | GBS_Bsp-like, Glyco_hydro_25 | Streptococcus milleri, Streptococcus sp. | | lysozyme activity [GO:0003796] |
| 83 | HxuA | A0A379IT80 | 2 | 88 | YDG | 9 | 10 | 0.82 | 0.9 | ESPR, Haemagg_act, YDG | Pseudomonas mendocina | | |
| 84 | UNK_Invasin | A0A380CN38 | 2 | 103 | Big_1, Invasin_D3 | 7 | 8 | 0.81 | 0.85 | Big_1, Big_9, Cadherin_5, CHU_C, DUF11, Invasin_D3 | Sphingobacterium spiritivorum | | |
